# Structural and immunological identification and antiviral infection experiment of the dominant cytotoxic T lymphocyte epitopes of the oncogenic Marek’s disease virus

**DOI:** 10.1101/2021.01.25.428198

**Authors:** Beibei Sun, Yawen Wang, Zhenbao Wang, Shuangshuang Lu, Chun Xia

**Author notes:** Address correspondence to Chun Xia,. Beibei Sun and Yawen Wang contributed equally to this work. Author order was determined on the basis of seniority.

## Abstract

Marek’s disease (MD) is a serious cancer caused by MDV in chickens, and it is also the first tumor disease that could be controlled by an attenuated vaccine in the world. However, the attenuated vaccine is able to inhibit only the formation of tumors but cannot prevent the infection of oncogenic viruses; instead, it leads to mutations and the emergence of a number of virulent strains. In this paper, the crystal structures of chicken MHC class I (pBF2*1501) molecules bound with 8-mer and 9-mer MDV peptides were solved. The results showed that the conformations of the 8-mer and 9-mer peptides in the antigen-binding groove (ABG) of pBF2*1501 are different; the ABG-8-mer is flat, and the ABG-9-mer has the “featured” M-type epitope morphology. Based on these results, multiple MDV epitopes were confirmed using the tetramer technique, and the immunoprotective effect of dominant epitope was confirmed using the protein adjuvant HSP108-N333 in BF2*1501-expressing chickens. The results showed that the two epitopes have obvious protective effects after a standard immunization program. After challenge, the mortality rate was only 20%, and the protective index (PI) reached 33% in the epitope group. The results verified that a single epitope could induce extremely strong specific antitumor T lymphocyte immunity. The results show that three key factors are crucial to obtain the best antitumor effect: accurate identification of dominant cytotoxic T lymphocyte (CTL) epitopes, efficient protein adjuvant and MHC matching. Meanwhile, the results also provided obvious advantages for the development of multiple epitope vaccines for tumor diseases.

**Importance:** Marek’s disease (MD) is the first tumor disease that can be controlled by an attenuated vaccine. However, the attenuated vaccine cannot prevent the infection of the oncogenic virus and instead leads to the emergence of a number of mutated virulent strains. First, the mechanism of the chicken BF2*1501 preferentially presents 9-mer peptide was clarified. Based on this result, multiple MDV epitopes were confirmed, and the dominant epitopes were identified. Subsequently, two epitope groups showed obvious protective effects after a standard immunization program, and the protective index (PI) of one of the epitope groups reached over 30%. The results showed that a single epitope could induce strong and efficient antitumor CTL immunity. Overall, three key factors are crucial to obtain the best antitumor effect: accurate identification of dominant epitopes, efficient protein adjuvant and MHC matching. Our results provide an obvious strategy for the development of multiple epitope vaccines in tumor diseases.

## Introduction

Marek’s disease (MD) is a serious cancer caused by Marek’s disease virus (MDV), which is an oncogenic alpha herpesvirus. In addition to causing a large number of deaths, it can also cause a low feed conversion rate, weight loss, egg production reduction, and immunosuppression, leading to secondary infection of other pathogens in chickens (1–4). MD was first described a hundred years ago, and after decades of research, scholars have made great efforts in understanding the genomics(5–7), serology (8), viral lifecycle (9, 10), pathogenesis of cancer (11, 12), vaccinology and virus mutation mechanism of MDV (13, 14). Therefore, a systematic knowledge of MD has been accumulated. With the emergence of mutant and very virulent MDV strains, immune-related failure events have occurred frequently, causing approximately 1-2 billion US dollars of economic losses annually in the world poultry industry. Since no specific antivirus antibody could be detected in chickens infected with MDV, the disease became the only tumor disease controlled by T cell immunity in animal viral diseases.

The MDV genome is a double-stranded DNA with a length of approximately 180 kb, which is composed of six regions: a unique long (U_L_) region, a unique short (U_S_) region, a terminal repeat long (T_RL_) region and an internal repeat long (I_RL_) region on both sides of the U_L_ region, and a terminal repeat short (T_RS_) and an inner short repeat (I_RS_) on both sides of the U_S_(15). MDV encodes more than 100 functional genes. The “Cornell model of MDV infection” explains the MDV lifecycle and related major functional genes(8, 16). By homology comparison with other herpesvirus-related genes, many functional genes have been identified (7, 17). It is worth noting that gB has been proven to be the main immunoprotective antigen, and its subunit vaccine can stimulate not only humoral immunity but also cellular immunity (1, 18). In addition to structural genes, seven MDV-specific genes were also identified: the oncoprotein gene Marek’s EcoRI-Q (Meq), 1.8 kb gene family, latent infection associated transcripts (LATs), viral telomerase RNA (vTR) gene, viral lipase (vLP) gene and IL8 gene (vIL8)(19). Among them, the most studied gene is the oncogenic Meq gene. Meq can not only form a homologous polymer (Meq/Meq) but also combine with c-JUN to form a heterologous polymer (Meq/JUN). Both of them contribute to the transformation of lymphocytes during MD in chickens(20). A Meq-null virus, rMd5ΔMeq, can replicate well in vitro but is not oncogenic, which supports that the Meq gene is closely related to MDV tumorigenesis(21). Overall, gB and Meq are immunoprotective genes, and both can induce specific cytotoxic T lymphocyte (CTL) immunity(22).

According to the different antigenicities, MDV can be classified into three serotypes: MDV1, MDV2, and MDV3. Through long-term evolution, the MDV1 lineage has been differentiated into a series of virulent strains, and all of the strains lead to tumors, immunosuppression and even high mortality in chickens; thus, MDV1 is the most serious serotype in chickens(23). Based on virulence, MDV1 has been divided into four pathogenic types, namely, mildly virulent MDV (mMDV), virulent MDV (vMDV), very virulent MDV (vvMDV), and very virulent plus MDV (vv^+^MDV) (24, 25). mMDV strains cause only mild neurological diseases; vMDV strains can cause lymphoma and have a mortality rate of up to 40%(24). vv^+^MDV strains often cause severe brain edema and acute death in nonvaccinated chickens and even tumor lesions in vaccinated chickens (26–29). To date, the virulence of MDV1 has been proposed to evolve towards stronger virulence, which poses a greater threat to the poultry industry (30).

Since the 1970s, veterinary scholars have first developed a live-attenuated vaccine and effectively controlled the outbreak of MD (31, 32). Currently, the most widely used and most effective vaccine is the live-attenuated CVI988/Rispens vaccine (32). Unfortunately, the vaccine is not perfect, and even though it can control the clinical development of tumors, it does not prevent viral infection and transmission. As a consequence, vaccinated chickens still become infected and shed the virus into the environment (33). In addition, the vaccine itself can also replicate and shed in chickens (2, 34). Thus, the live-attenuated vaccine can reduce mortality and prolong the survival time of infected chickens, but it does not significantly reduce the virus excretion rate (35). Decades of using such an “imperfect” MDV vaccine inevitably led to the extensive evolution and spread of MDV1 in the field, resulting in the emergence of more virulent mutants (36). With the use of an attenuated vaccine, the virulence of MDV1 is also evolving while protecting chickens from MDV1-induced tumor symptoms (14). Since the vaccine was first used in 1969, the virulence of MDV1 has evolved from mMDV to vv+MDV(37). With the emergence of new and more virulent strains, the CVI988/Rispens vaccine has gradually failed to provide complete protection in commercial chicken flocks, which often leads to outbreaks of MD (38, 39). Therefore, there is a need for exploring and developing safer and more effective vaccines for MD.

Genetically engineered vaccines have become a way to develop MD vaccines. Recombinant fowl poxviruses (rFPVs) expressing the MDV1 gB gene effectively protect against challenge with the Md5 strain (40). The Meq-null virus rMd5Δmeq provides superior protection than CVI988/Rispens against challenge with the 648A strain (41). The GX0101Δmeq deletion strain SC9-1 constructed by bacterial artificial chromosome (BAC) technology can induce better protection than CVI988/Rispens in vivo and can cause a loss of tumorigenicity(42). In recent years, the CRISPR/Cas9 system has been successfully used to construct Meq gene deletion strains(43). A DNA vaccine containing the BAC20 clone of MDV1 has also been proven to be resistant to vMDV to some extent (17). Unlike genetically engineered vaccines, epitope vaccines have not been reported, possibly because it is not known how to establish an effective system in epitope vaccines to identify epitopes and how to break the restriction between major histocompatibility complex class I/II (MHC-I/II) and T-cell receptor (TCR) molecules.

Epitope vaccines are usually made of epitopes and effective adjuvants. Because of its high biological safety, good immunogenicity, easy transportation and preservation, it has become a popular research topic in the prevention and control of tumors and infectious diseases (44, 45). The CTL epitope vaccine derived from human Wilms’ tumor gene 1 (WT1) has been proven to induce effective immune responses in patients with various hematological tumors and various types of solid tumors (46). The human leukocyte antigen (HLA)-restricted gp100-in4 peptide vaccination can induce antigen-specific T cell responses in multiple patients (47). The E75 vaccine derived from the HER2/neu protein can induce effective specific immunity against breast cancer in vivo(48). The multiepitope vaccine derived from the HIV-1 integrase protein has been proven to have antigenicity and can induce effective immunity to HIV-1 (49). An H5N1-based matrix protein 2 ectodomain tetrameric peptide vaccine provides cross-protection against lethal infection from the H7N9 influenza virus (50). In the prevention and control of infectious diseases in livestock, B- and T helper (Th)-cell epitope vaccines derived from foot-and-mouth disease virus (FMDV) showed 60% protection against serotype-A FMDV infection in cattle (51). These results bring more hope to epitope vaccines in the prevention and control of cancers and infectious diseases.

The survival and proliferation of MDV are strictly dependent on host cells, and the CTL immune response plays a critical role in the process of eliminating MDV1-infected cells(22). In this paper, the crystal structures of peptide/MHC-I (pMHC-I aka pBF2*1501) molecules bound with 8-mer and 9-mer peptides were first solved. The results showed that the conformations of the 9-mer peptide in the antigen-binding groove (ABG) of pBF2*1501 has the “featured” the M-type epitope morphology. Based on this antigen presentation mechanism, multiple MDV epitopes were confirmed using the tetramer technique. The immunoprotective effect of two dominant epitope peptides was proven in BF2*1501 specific-pathogen-free (SPF) chickens. The results showed that the two epitopes have obvious protective effects after an immunization program. For a single epitope, the mortality rate was only 20%, and the PI reached 33%. The results show that three key factors are crucial to obtain the best antitumor effect: accurate identification of dominant CTL epitopes, efficient protein adjuvant and MHC matching. The results also provided obvious advantages for the development of multiple epitope vaccines in animals and humans.

## Results

### Structural mechanism of the BF2*1501 presenting peptide profile

The crystal structures of pBF2*1501_-meq-RY0801_ and pBF2*1501_-meq-RY0901_ were determined (Table 1, Fig. 1A and 1B). The two structures were similar, and the root mean square deviation (RMSD) was 0.227 (Fig. 1C). Superposition of pBF2*1501_-meq-RY0901_ with the known pMHC-I complexes of representative species (pHLA-Cw4, PDB ID: 1IM9; pMamu-A*01, PDB ID: 1ZVS; pBF2*2101, PDB ID: 3BEW; pBF2*0401, PDB ID: 4E0R; pXela-UAAg, PDB ID: 6A2B; and Ctid-UAA*0102, PDB ID: 5H5Z) showed that the structure of pMHC-I molecules within chickens is basically regular. pBF2*1501 is similar to pBF2*2101 and pBF2*0401, with RMSDs of 0.484 and 0.757, respectively, whereas the RMSD of pBF2*1501 with monkey pMamu-A*01 is 1.685 (Fig. 1C). The results of aa sequence alignment of chicken MHC-I showed that the identity of BF2*1501 with BF2*2101 and BF2*0401 was 90.7% and 87.4%, respectively (Fig. 2), the sequence differences were mainly manifested in the α1 and α2 regions.

**Figure 1.**
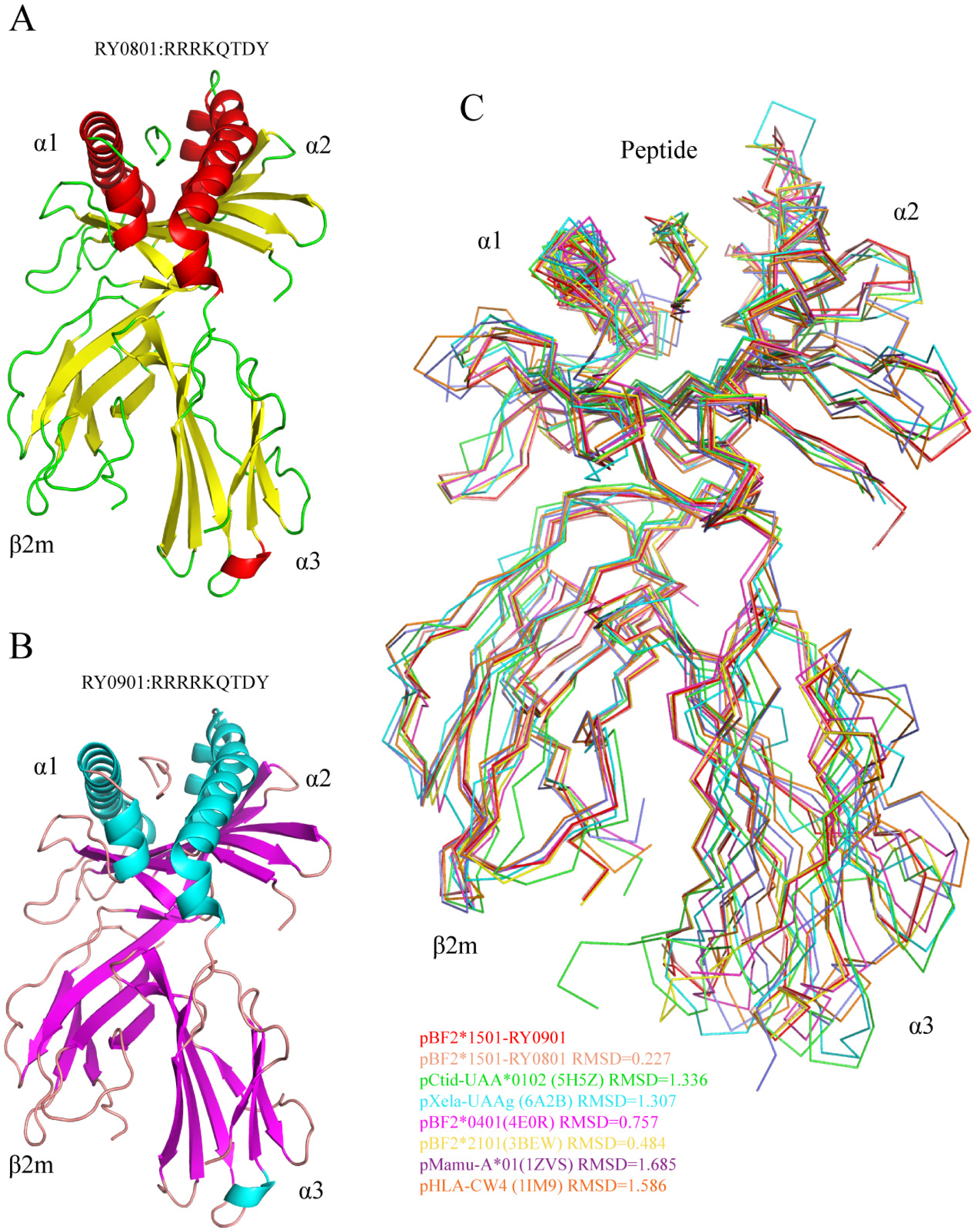
Structural overview of the pBF2*1501 complex. (A) The structure of pBF2*1501 with the 8-mer Meq-RY0801, in which the α-helix is shown in red, the β-sheet is shown in yellow, and the loop is shown in green. (B) The structure of pBF2*1501 with the 9-mer Meq-RY0901, in which the α-helix is shown in cyan, the β-sheet is shown in magenta, and the loop is shown in pink. (C) The superposition of pBF2*1501_-meq-RY0901_ with the known pMHC-I structures of representative species (HLA-CW4, PDB ID: 1IM9; pMamu-A*01, PDB ID: 1ZVS; pBF2*2101, PDB ID: 3BEW; pBF2*0401, PDB ID: 4E0R; pXela-UAAg, PDB ID: 6A2B; and pCtid-UAA*0102, PDB ID: 5H5Z). The RMSD values are shown.

**Figure 2.**
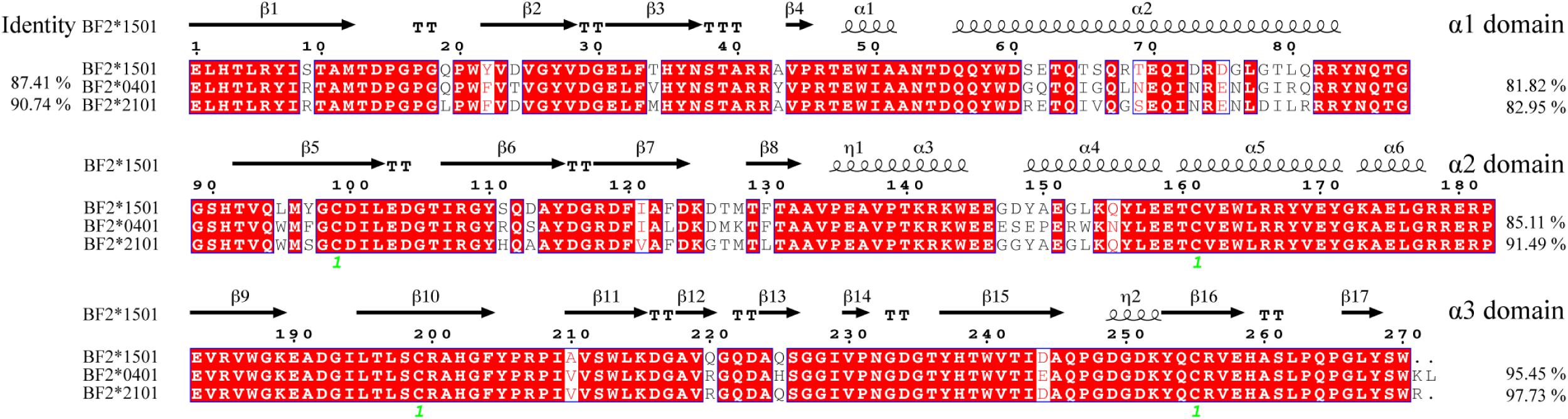
Structure-based amino acid sequence alignment of chicken MHC-I molecules among BF2*1501, BF2*0401 and BF2*2101. The total amino acid identities among BF2*1501, BF2*0401 and BF2*2101 are shown at the beginning of each sequence, while individual identities of the α1, α2 and α3 domains are listed at the end of each sequence. The cysteine residues forming the disulfide bond are marked by a green “1”. Conserved residues are highlighted in red.

**Table 1.** X-ray diffraction data collection and refinement statistics

| Parameter | BF2*1502-RY0801 | BF2*1501-RY0901 |
| --- | --- | --- |
| **Data processing** |  |  |
| Space group | P3<sub>1</sub>21 | P3<sub>1</sub>21 |
| Cell parameters (Å) | $a=125.07$ $b=125.07$<br>$c=80.64$ | $a=125.14$, $b=125.14$, $c=80.87$ |
| | $\alpha=90.0$, $\beta=90.0$, $\gamma=120.0$ | $\alpha=90.0$, $\beta=90.0$, $\gamma=120.0$ |
| Volume of the unit cell(Å<sup>3</sup>) | 1092494.1 | 1096701.6 |
| Resolution range (Å) | 50.0–2.85(2.95–2.85)<sup>a</sup> | 50.0–2.6(2.698–2.6)<sup>a</sup> |
| Total No. of reflections | 83861 | 205783 |
| No. of unique reflections | 10340 | 24077 |
| Completeness (%) | 99.4(100.0) | 100.0 (100.0) |
| Average redundancy | 4.9(4.9) | 9.0(9.1) |
| $R_{\text{merge}}$ (%)<sup>b</sup> | 12.4(56.6) | 11.7 (54.0) |
| $I/\sigma$ | 14.1(3.1) | 22.1(5.2) |
| $R_{\text{work}} / R_{\text{free}}$<sup>c</sup> | 0.177/0.227 | 0.172/0.225 |

<sup>a</sup> Values in parentheses are for the highest-resolution shell.
<sup>b</sup> $R_{\text{merge}} = \sum_{hkl} \sum_i |I_i(hkl) - \langle I(hkl) \rangle| / \sum_{hkl} \sum_i I_i(hkl)$ , where $I_i(hkl)$ is the observed intensity, and $\langle I(hkl) \rangle$ is the average intensity from multiple measurements.
<sup>c</sup> $R = \sum_{hkl} ||F_{\text{obs}}| - k|F_{\text{calc}}| / \sum_{hkl} |F_{\text{obs}}|$ , where $R_{\text{free}}$ is calculated for a randomly chosen 5% of reflections, and $R_{\text{work}}$ is calculated for the remaining 95% of reflections used for structure refinement.

In the structures of pBF2*1501_-meq-RY0801_ and pBF2*1501_-meq-RY0901_, the orientations of the 8-mer and 9-mer peptides in the ABG were basically the same (Fig. 3A). P2 and P8/9 are the main anchor residues, which bind to the B and F pockets, respectively, to stabilize the conformation of the ABG of pMHC-I. What is interesting is the difference in the conformation of the two peptides in the ABG. Compared with the “flat” conformation of Meq-RY0801, the middle part of Meq-RY0901 forms a higher protrusion, which has a more “featured” M-type epitope morphology (Fig. 3A).

**Figure 3.**
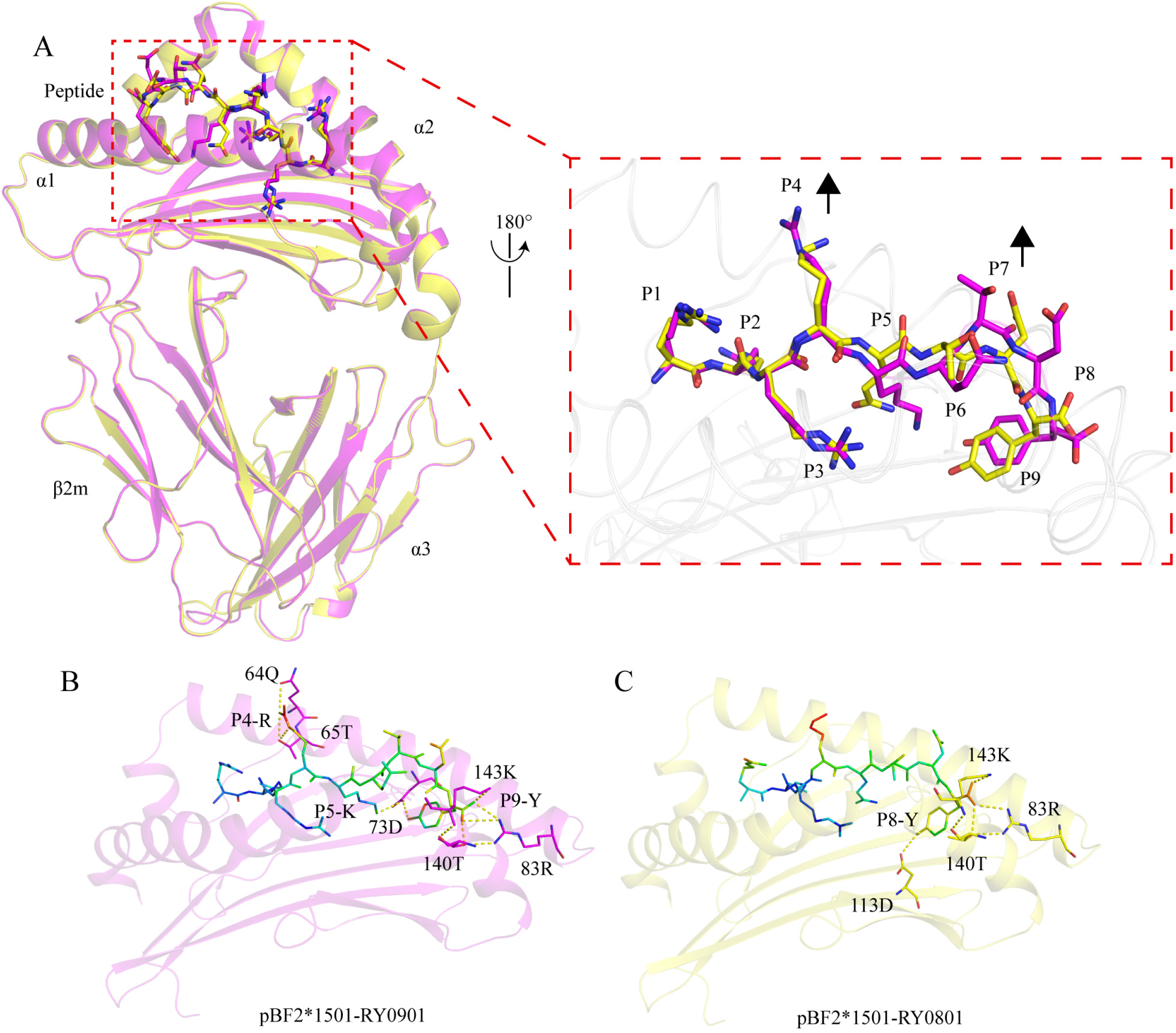
The structure of pBF2*1501_-meq-RY0901_ has the “featured” M-type conformation. (A) Comparison of the pBF2*1501_-meq-RY0901_ and pBF2*1501_-meq-RY0801_ structures; pBF2*1501_-meq-RY0901_ is shown in magenta, and pBF2*1501_-meq-RY0901_ is shown in yellow. The magnified inset on the right shows the presentation conformation of the 8-mer-RY0801 and the 9-mer-RY0901. The position of the 9-mer peptide is higher than that of the 8-mer peptide at P4 and P7-8. (B and C) The key hydrogen bonds in the formation of the RY0901 and RY0801 conformations, respectively. The hydrogen bonds are represented by yellow dotted lines. The corresponding residues are labeled with amino acid abbreviations and primary sequence numbers.

Meq-RY0901 P4-Arg forms a hydrogen bond with Gln64 and Thr65 in the ABG, P5-Lys forms a hydrogen bond with Asp73, and P9-Tyr forms a hydrogen bond with Arg83 and Lys143. These forces make the 9-mer peptide have more “featured” characteristics than the 8-mer peptide (Fig. 3B). The hydrogen bonds formed by the 8-mer-RY0801 P8-Tyr and Arg83, Asp113, Thr140, and Lys143 together form an inward and downward force, especially Asp113 at the bottom of the ABG, and Asp113-P8 makes the C-terminus of the 8-mer-RY0801 insert deeper into the ABG, resulting in a flatter conformation (Fig. 3C). At the same time, we also found that the hydrogen bond network formed between 9-mer-RY0901 and the ABG is significantly stronger than that of 8-mer-RY0801. Meq-RY0901 P2-Arg can form 4 pairs of salt bridges with Asp24, Thr34, and Glu62 in the ABG, while RY0801 P2-Arg has only three salt bridges in the ABG (S1 Table). In the P4-P6 regions, Meq-RY0801 lacks intermolecular hydrogen bonding, while Meq-RY0901 forms a relatively stable hydrogen bond network (S1 Table). In addition, the ASA values of 9-mer-RY0901 and 8-mer-RY0801 are 1635.97 Å^2^ and 1515.61 Å^2^, respectively; the BSA values are 1171.03 Å^2^ and 1167.84 Å^2^, respectively (S2 Table). The ASA of RY0901 is 120.36 Å^2^ more than that of Meq-RY0801, which is mainly concentrated at the positions of P4-P8/7. The ASA values of Meq-RY0901 and Meq-RY0801 in P4-P8/7 are 716.46 Å^2^ and 526.22 Å^2^, respectively (S2 Table), which is a difference of 140.40 Å^2^. Obviously, the part of 9-mer-RY0901 that is exposed is more than the part of 8-mer-RY0801 and is concentrated in the TCR recognition site(52), suggesting that nonapeptides are more easily recognized by the TCR.

### Structural insights into the chicken BF2*1501 haplotype on the resistance to MDV

Chicken MHC-I is not only directly related to epitope vaccine responsiveness but also related to disease resistance. B21 chickens are genetically resistant to MD, while B4 and B15 chickens are genetically susceptible or have extremely low resistance to MD(53). Previous reports found that the B21 haplotype chicken MHC-I molecule BF2*2101 has a relatively wide ABG (Fig. 4A); therefore, BF2*2101 does not have a fixed anchor sequence when binding peptides, and the charged properties of the pocket are uneven. Arg9 in the ABG can mediate charge transfer, enabling BF2*2101 to bind more antigen peptides of different properties, thereby activating a large number of CTLs to mediate genetic resistance to MD(54). Although BF2*1501 also has a relatively wide ABG (Fig. 4B), the C, D and E pocket boundaries are clear, and the volume is relatively narrow (Fig. 4C), which reduces the probability of BF2*1501 presenting long peptides or long side chain peptides. As the anchor pocket, the B and F pockets are both negatively charged, which determines that the P2 and P8/9 positions of the binding peptides cannot be negatively charged residues, which further restricts the kinetics of binding epitope peptides (Fig. 4D and 4E). This is the key knowledge point for designing epitope vaccines in B15 chickens.

**Figure 4.**
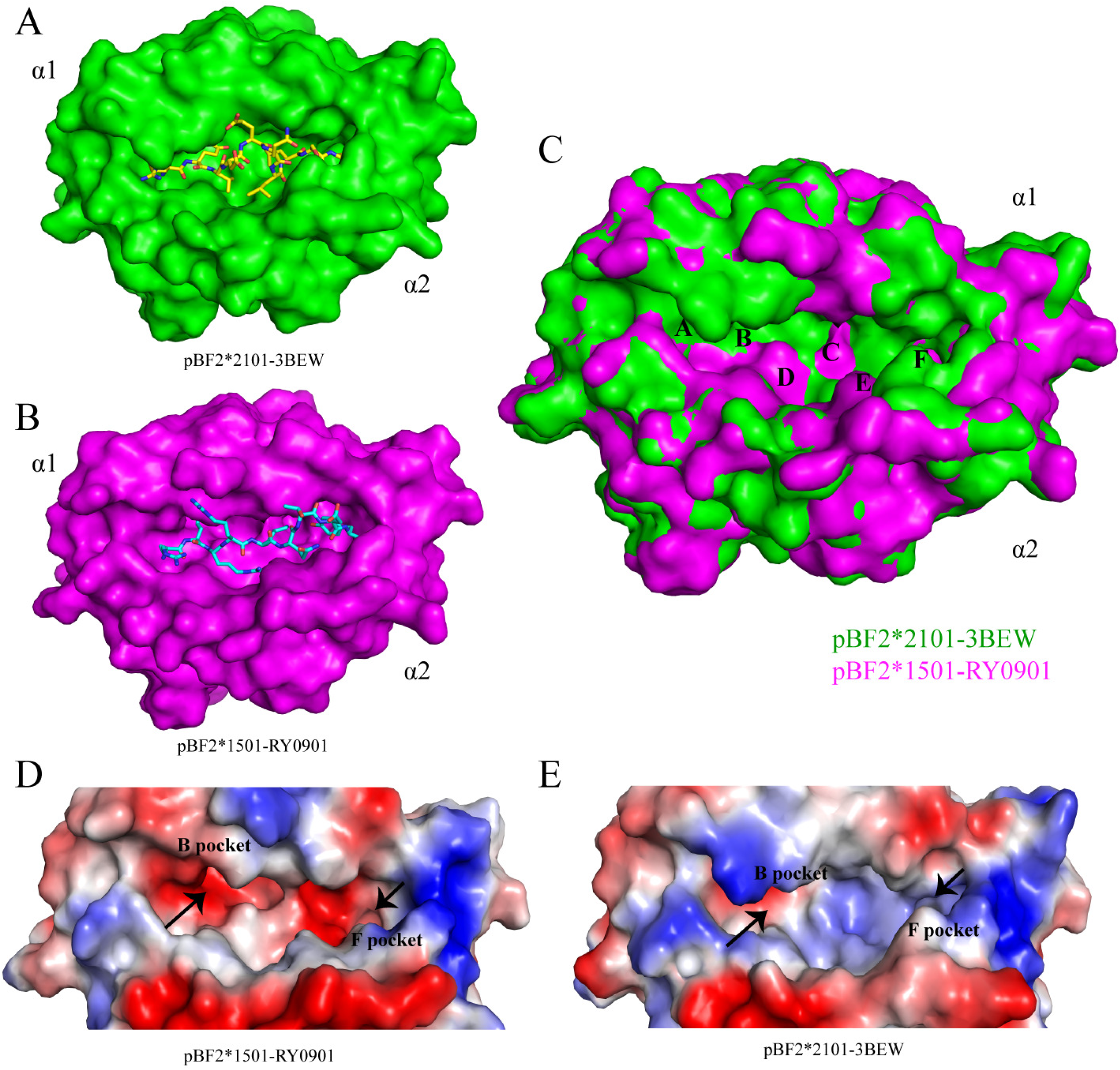
Restriction of the pBF2*1501 pocket in presenting peptides. (A and B) The pocket conformations of pBF2*2101 (green) and pBF2*1501 (magenta) are shown in surface representation. (C) Superposition of the α1 and α2 domains of pBF2*2101 (green) with pBF2*1501 (magenta) are shown in surface representation. We found that the C, D, and E pockets of pBF2*1501 are smaller than those of pBF2*2101. (D and E) The pocket charge distributions for pBF2*1501 and pBF2*2101 (blue, positively charged; red, negatively charged; and white, nonpolar). The B and F pockets of pBF2*1501 have a strong negative charge, while the B and F pockets of pBF2*2101 has an uneven charge.

### Primary selection and identification of dominant MDV epitopes

The peptide-binding motif of the BF2*1501 allele as X-R-X-X-X-X-(X/XX)-Y has been determined(55). Based on the motif and using the epitope prediction tool (CBS Prediction Servers), whole MDV1 proteins were scanned, and 15 peptides were obtained (S3 Table). Among them, 13 nonapeptides were refolded in vitro with BF2*1501 and chicken β2m and finally purified by volume exclusion chromatography and ion exchange chromatography except for GB0901 and GB0904. Subsequently, BF2*1501 tetramers loaded with GB0902, GB0903, GB0905 and Meq-RY0903 were constructed (Fig. 5). After B15 chickens were immunized with the live-attenuated CVI988/Rispens vaccine, peptide-specific CD8^+^ T cells (FITC-labeled anti-chicken CD8^+^) and PE-labeled tetramer (Tet^+^) double-positive CTLs were detected by constructing specific tetramers (Fig. 6A). The results showed that the proportion of CD8^+^Tet^+^ cells in the total number of lymphocytes in the control group was 0.07-0.17%, and the distribution of lymphocytes was relatively scattered, which was determined to be nonspecific T cells (Fig. 6B). The lymphocytes in the immune group were more concentrated and were specific T cells. Among them, CD8^+^Tet^+^ GB0902 cells accounted for 2.77-3.88%, CD8^+^Tet^+^ GB0903 cells accounted for 1.77-2.19%, CD8^+^Tet^+^ GB0905 cells accounted for 1.29-1.57%, and CD8^+^Tet^+^ Meq-RY0903 cells accounted for 0.42-0.50% (Fig. 6B). Further analysis of significant differences in the data between the groups showed that the proportions of CD8^+^Tet^+^ GB0902 T cells, CD8^+^Tet^+^ GB0903 T cells, CD8^+^Tet^+^ GB0905 T cells and CD8^+^Tet^+^ Meq-RY0903 T cells in the sample group were significantly higher than those in the control group, with P values of 0.0007, 0.0001, 0.0001, and 0.0009 (P<0.05 indicates a significant difference) (Fig. 6C). The GB0902, GB0903, GB0905 and Meq-RY0903 peptides could be presented by BF2*1501 on antigen-presenting cells (APCs) and recognized by specific CTLs though pBF2*1501 in vivo. However, the proportion of double-positive CD8 T cells stimulated by GB0902 and GB0903 was significantly higher than that stimulated by GB0905 and RY0903 (P<0.05). Therefore, GB0902 and GB0903 were identified as MDV1-dominant CTL epitopes for further experiments.

**Figure 5.**
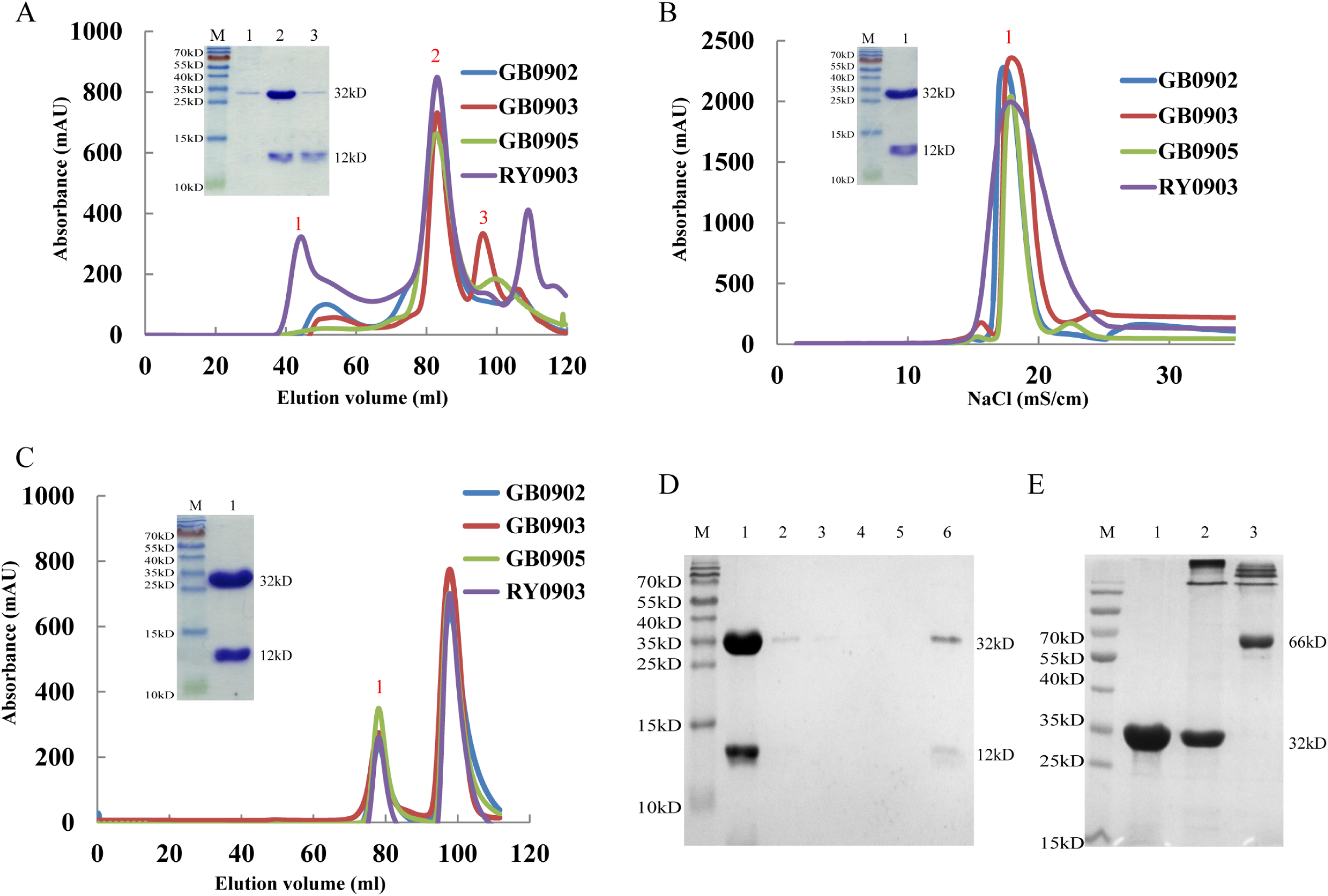
Production of the BF2*1501 tetramers. (A) Gel filtration chromatograms of the pBF2*1501-BSP complex with the four MDV peptides. Peaks 1, 2 and 3 represent the aggregated HC (32 kDa), the correctly refolded pBF2*1501-BSP complex (45 kDa), and the extra chicken β2m (12 kDa), respectively. (B) Ion-exchange chromatogram of the pBF2*1501-BSP complex with the four MDV peptides. Peak 1 represents the correctly refolded pBF2*1501-BSP complex (45 kDa) at a NaCl concentration of 15–20 mS/cm. (C) Gel filtration chromatograms of the biotinylated pBF2*1501-BSP complex with the MDV peptides. Peak 1 represents the correctly biotinylated pBF2*1501 complex (45 kDa). (D) SDS-PAGE analysis of the efficiency of pBF2*1501-BSP biotinylation. The biotinylated pBF2*1501-BSP was mixed with streptavidin MagneSpheres. Lane 1 represents the supernatant from biotinylated pBF2*1501-BSP that reacted with streptavidin MagneSpheres. Lanes 2, 3, 4 and 5 contain the supernatants from the first, second, third and fourth washings of the streptavidin MagneSpheres, respectively. Lane 6 contains the supernatant of streptavidin MagneSpheres boiled after washing the sample four times. (E) SDS-PAGE analysis of the efficiency of the tetramers. Biotinylated pBF2*1501-BSP was mixed with PE-labeled streptavidin and filtered with a 100 kDa Millipore filter. Lane 1, biotinylated pBF2*1501-BSP monomer; lane 2, pBF2*1501-BSP tetramer >100 kDa; and lane 3, PE-labeled streptavidin.

**Figure 6.**
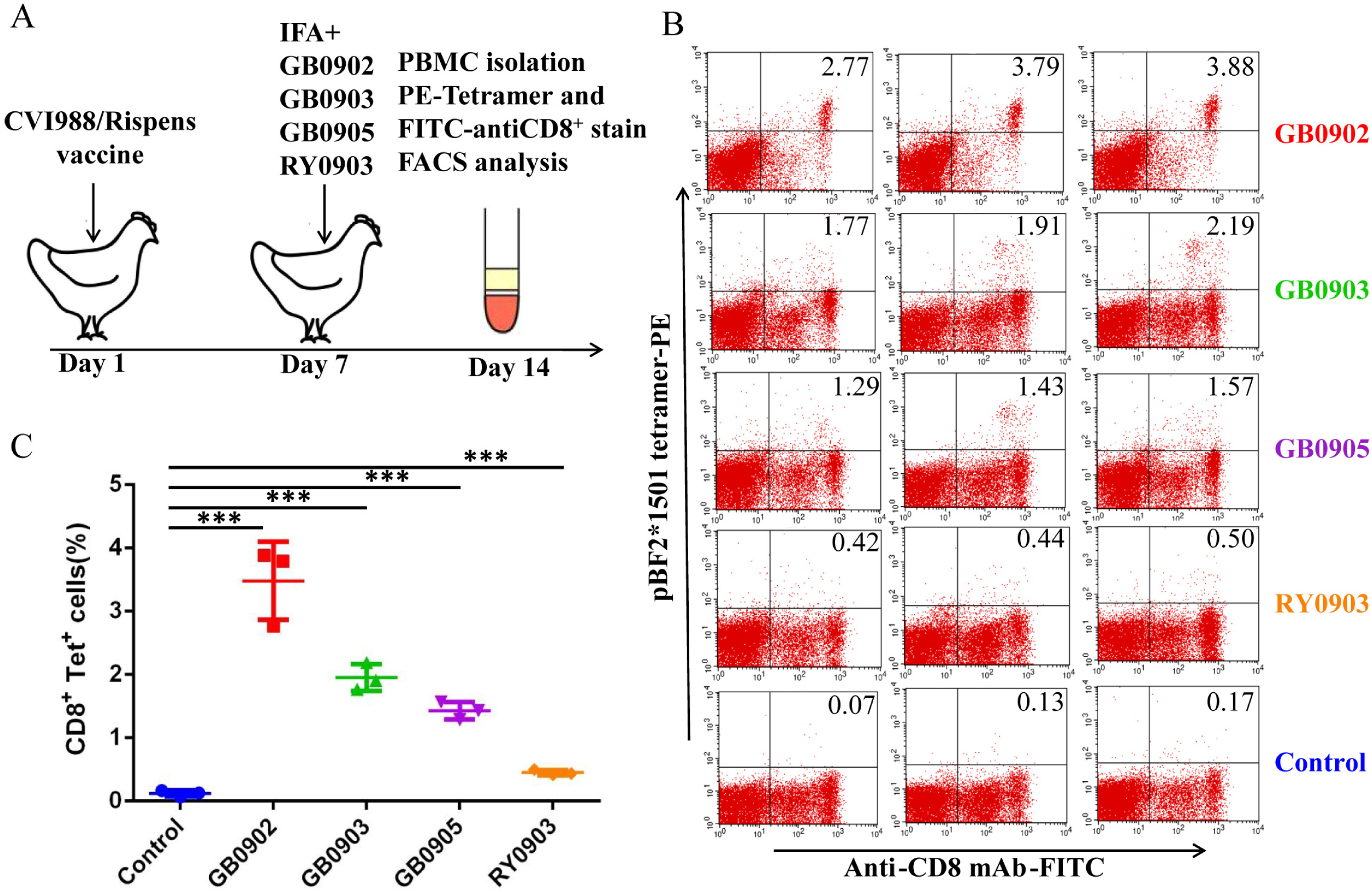
Identification of the functional MDV CTL epitopes. (A) The strategy of immunizing SPF B15 chickens with the commercially available CVI988/Rispens vaccine and peptides to detect CD8+ T cells specific for the GB0902, GB0903, GB0905 and Meq-RY0903 epitopes. (B) Flow cytometry was used to analyze CD8+ and pBF2*1501 Tet+ cells. Data are shown as pseudocolor plots. (C) Statistical data were analyzed by GraphPad Prism. The percentages of GB0902, GB0903, GB0905 and Meq-RY0903 epitope-specific CD8+ T cells were significantly higher in the vaccine-immunized group than in the control group (P < 0.01). *** means P < 0.001 by the unpaired Student’s t-test.

### Human Gp96 analog chicken HSP108-N333 protein adjuvant can increase the immune effect of epitopes by more than 9 times

The DNA fragment of HSP108-N333 was amplified by RT-PCR from chicken liver (Fig. 7A), and HSP108-N333 protein was obtained by prokaryotic expression (Fig. 7B). The purified HSP108-N333 protein (100 µg) was combined with the CTL epitope peptide GB0903 (100 µg) in low salt binding buffer, and chickens were immunized with the emulsified peptide vaccine. PBMCs were collected and stained with the PE-labeled tetramer and FITC-labeled anti-chicken CD8 monoclonal antibody as mentioned above. The ratio of double-positive CTLs was analyzed by flow cytometry. The results showed that the ratio of double-positive T cells in the control group was 0%, the ratio of double-positive T cells in the group immunized with only the GB0903 epitope was 0.8%, and the ratio of double-positive T cells in the group immunized with the GB0903 epitope plus the HSP108-N333 adjuvant was 7.5% (Fig. 7C). The results showed that the HSP108-N333 protein adjuvant can increase the immunization effect of epitopes by more than 9.2 times.

**Figure 7.**
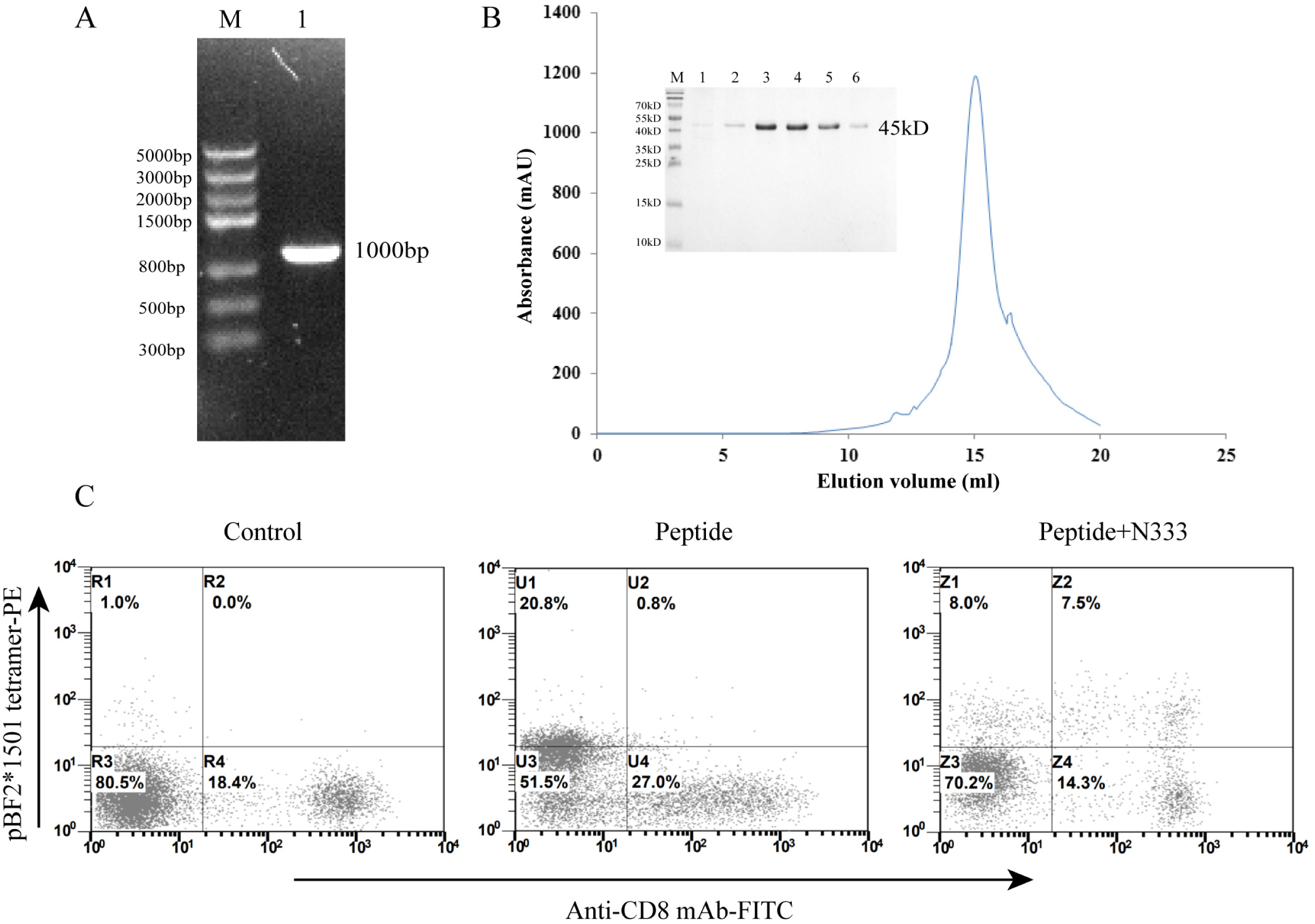
Expression, purification and effect analysis of N333 protein immunoadjuvant. (A) The PCR amplification result of the HSP108-N333 gene showed that the target band was approximately 1000 bp. (B) N333 protein purification results. The target protein was obtained at an approximately the 15-ml peak, with a molecular size of 45 kDa. Lanes 1-6 represent six tubes of samples collected around the peak. (C) The flow cytometry results of the control group, peptide-immunized group, peptide-immunized group and N333 protein adjuvant-immunized group from left to right. The ratio of double-positive CTLs was 0.0% in the negative control group, 0.8% in the peptide-immunized group, and 7.5% in the peptide and N333 protein adjuvant-immunized groups.

### Clinical symptoms and necropsy observation

The chickens in each group were observed for 42 days after immunization and challenge. The chickens in the control group grew normally, were in good spirits and were of uniform size, and there were no deaths during the whole experiment. The chickens in the challenge group had slow growth, were short and thin, were disheveled and had dull feathers, and four chickens died during the experiment; the earliest death occurred on the 19th day (31 days of age) after the MDV challenge, and the liver, spleen and kidney showed white tumor nodules. Some chickens appeared symptoms of glandular stomach, kidney enlargement and liver diffuse bleeding (Fig. 8A). In the GB0902 epitope group, 1 chicken died on the 30th day (42 days old) after the MDV challenge, and 1 chicken died on the 38 days (50 days old) after the MDV challenge. In the GB0903 epitope group, 2 chickens died on the 35th day (47 days of age) after the MDV challenge, and 1 chicken died on the 39th day (51 days of age) after the MDV challenge. Finally, all surviving chickens were sacrificed and dissected on the 42nd day (54 days of age). The results showed that there were no tumors in the control group. In the challenge group, multiple tumors were found in the liver, kidney and spleen, and were occasionally found in the heart. In addition, two chickens in the GB0902 epitope group had tumors in the liver and spleen respectively. Two chickens in the GB0903 epitope group had tumors in the liver and kidney respectively. (Fig. 8A).

**Figure 8.**
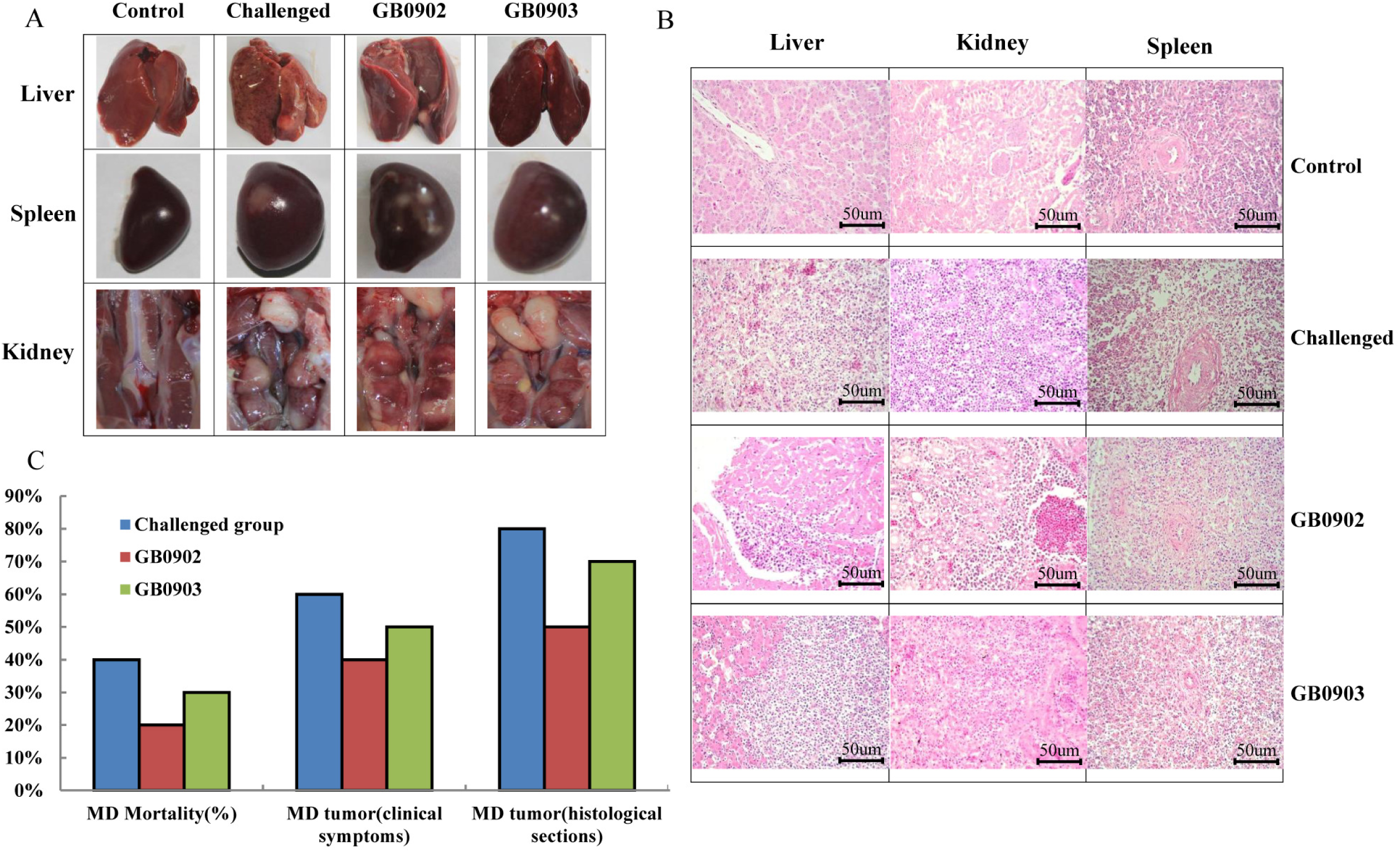
Evaluation of the immunogenicity of dominant epitopes. (A) Clinical symptoms after challenge with the vvMDV J-1 strain. MD-specific tumors mainly occurred in the liver, spleen and kidney. (B) Results of Histological section. MD-specific pathological features were observed in the liver, kidney and spleen. (C) Results of MD mortality, MD tumor (clinical symptoms) and MD tumor (histological section) in the challenge, GB0902, and GB0903 groups.

### Neoplastic histopathologic examination

To more objectively evaluate the protective effects of the GB0902 and GB0903 epitopes, the liver, spleen and kidney of all experimental chickens were used for paraffin sections, and hematoxylin-eosin staining at the same magnification (100x) was used for pathological observation (Fig. 8B). The organs of the control group were normal, with intact hepatocytes; the central artery was seen in the spleen; normal renal tubules and glomeruli were seen in the kidney. In the challenge group, a large number of tumor cells infiltrated into the liver to form tumor foci, and the intact liver cells were occupied by tumor cells; the tumor was not obvious in the spleen because of its own lymphocyte composition, but tumor cells could still be observed. The normal renal tubules were destroyed, and a large amount of neovascularization stimulated by tumor cells was found. The liver, spleen and kidney of some chickens in the GB0902 and GB0903 epitope groups showed different degrees of tumor cell infiltration (Fig. 8B). However, overall, the damage was more obvious in the challenge group than in the GB0902 and GB0903 epitope groups. The statistical results showed that the control group did not have MD-specific pathological changes; the challenge group showed different degrees of MD histological lesions in 8/10 sections, of which 4/8 had more than 2 tissue lesions. In the GB0902 epitope group, 5/10 sections showed different degrees of tumor lesions, of which 2/5 had more than 2 tissue lesions. In the GB0903 epitope group, 7/10 sections showed different degrees of tumor lesions, of which 3/7 had more than 2 tissue lesions. From the perspective of pathological changes, compared with that in the challenge group, the proportion of tumors in the GB0902 and GB0903 epitope groups was significantly reduced.

### Epitope immune effect

According to the clinical symptoms and necropsy results, the mortality rate of the challenge group was 40%, the tumor incidence rate was 60%, and the DI was 1.33. In the GB0902 epitope group, the mortality rate was 20%, the tumor incidence was 40%, the PI reached 33%, and the DI decreased to 1. In the GB0903 epitope group, the mortality rate was 30%, the tumor incidence rate was 50%, the PI reached 17%, and the DI decreased to 1. The control group had no deaths or tumors (Fig. 8C, Table 2). Tissue section results showed that the tumor incidence rate in the challenge group was 80%, and the DI was 1.5. However, the tumor incidence rate in the GB0902 epitope group was 50%, the PI reached 37.5%, and the DI decreased to 1.4. The tumor incidence rate in the GB0903 epitope group was 70%, the PI reached 12.5%, and the DI decreased to 1.43 (Fig. 8C, Table 3). In conclusion, the GB0902 and GB0903 epitope groups have obvious immunoprotective effects against vvMDV strain challenge.

**Table 2.** Protective efficacy of GB0902 and GB0903 against vvMDV J-1 chanllenge in SPF B15 chickens based on clinical symptoms

| Vaccine | Challenged with<br>J-1 at days | MD<br>Mortality (%) | MD<br>Tumor (%)<sup>a</sup> | PI (%) | DI |
| --- | --- | --- | --- | --- | --- |
| PBS | - | 10/0 (0) | 10/0 (0) | - | - |
| PBS | 12 | 10/4 (40) | 10/6 (60) | - | 1.33 |
| GB0902 | 12 | 10/2 (20) | 10/4 (40) | 33 | 1 |
| GB0903 | 12 | 10/3 (30) | 10/5 (50) | 17 | 1 |

<sup>a</sup>Based on clinical symptoms during the experiment

**Table 3.**
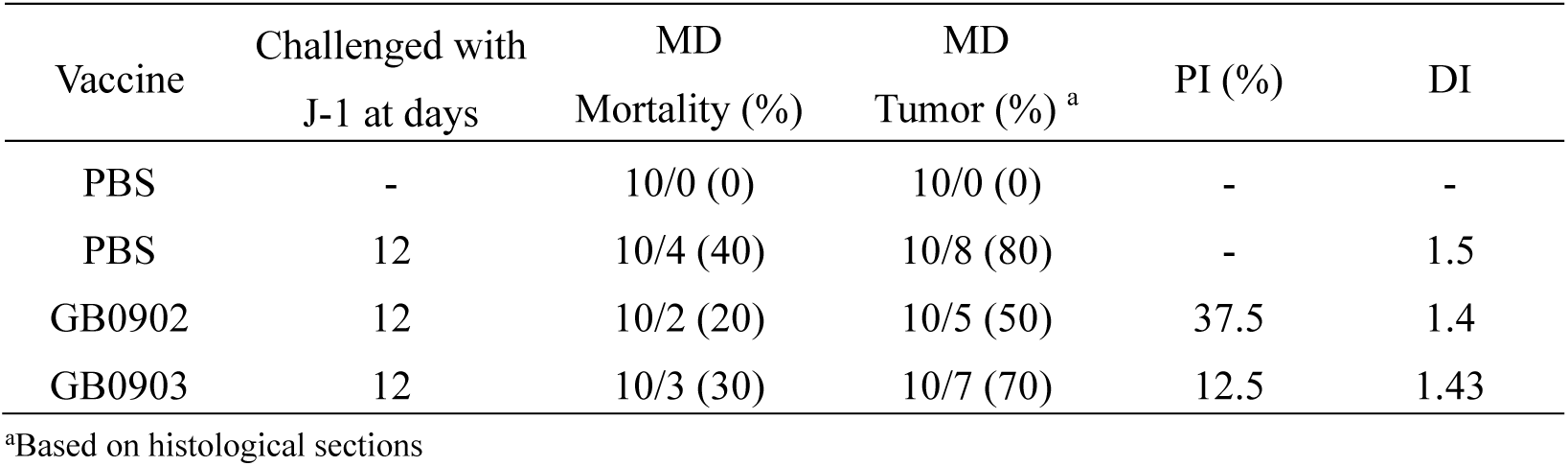
Protective efficacy of GB0902 and GB0903 against vvMDV J-1 chanllenge in SPF B15 chickens based on histological sections

### Determining the antigen presentation profile of the B15 haplotype and searching for candidate epitopes in MDV strains

To explore the antigen presentation profile by the B15 haplotype, BF2*1501, β2m and the random nonapeptide library Ran_9X were refolded in vitro, and the pBF2*1501 complexes were collected (Fig. 9A and 9B). The peptide sequence was determined by liquid chromatography–tandem mass spectrometry (LC-MS/MS), and 739 reliable nonapeptides were detected (S4 Table); the results are displayed in a map form by the WebLogo online tool (Fig. 9C). From the results, Arg, Lys and Leu often appear in the P1 position, Arg and Leu often appear in the P2 position, and Arg and Tyr frequently appear in the P9 position. These results are basically consistent with the results of previous studies(55); however, these results have been expanded in this study.

**Figure 9.**
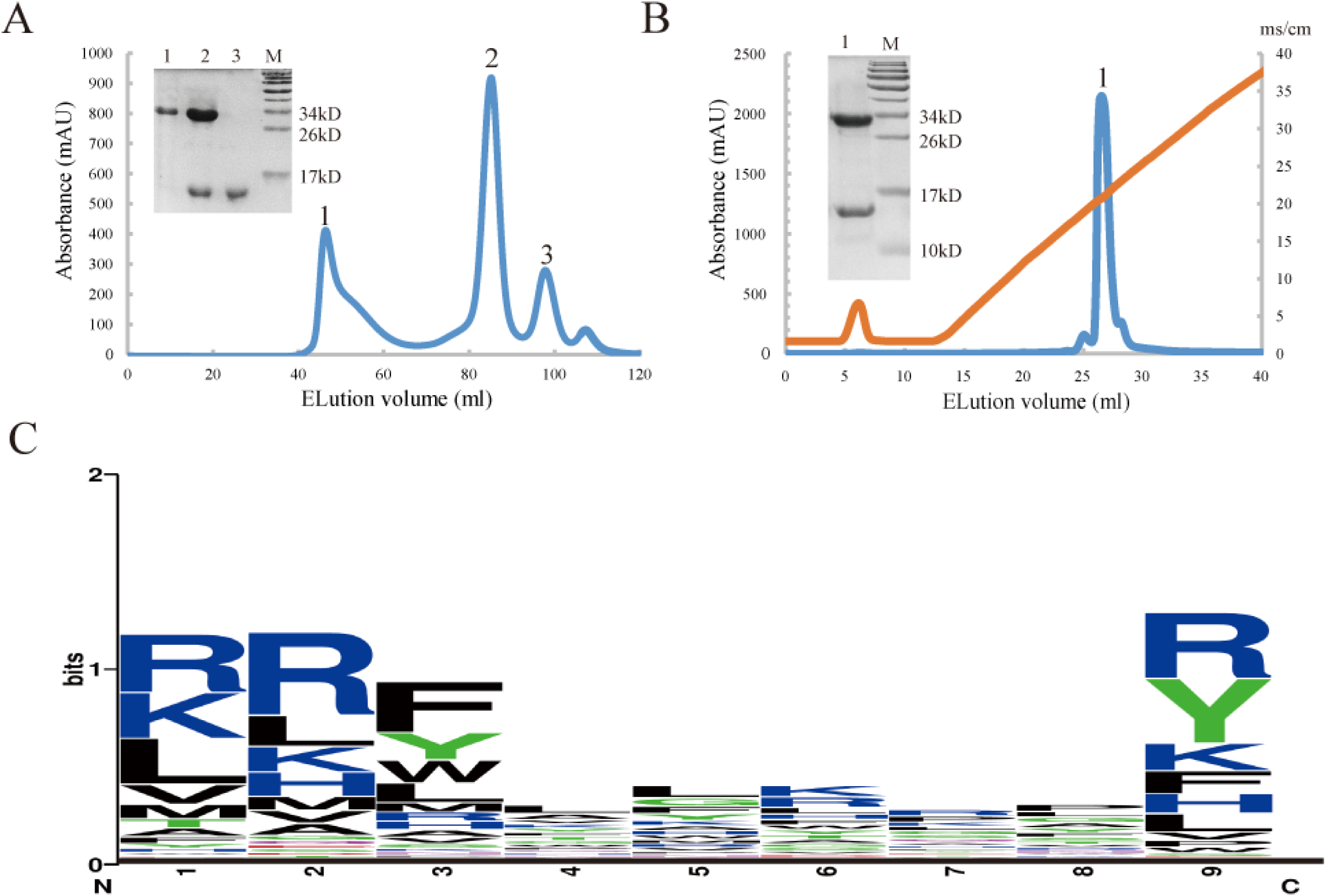
The peptide-binding motif for BF2*1501 identified by a random nonapeptide library. (A) Gel filtration chromatograms of the random nonapeptide library refolding with BF2*1501 and chicken β2m in vitro. Peaks 1, 2 and 3 represent the aggregated HC (32 kDa), the correctly refolded pBF2*1501 complex (45 kDa), and the extra β2m (12 kDa), respectively. The abscissa shows the peak volume (ml), whereas the ordinate represents the UV intensity (mAU). The insets show the SDS-PAGE results of the peaks that are labeled on the curve. Lane M contains molecular mass marker. (B) Ion-exchange chromatogram of the random nonapeptide library refolding with BF2*1501 and chicken β2m in vitro. (C) The sequence showing the amino acid weighting probabilities at every position of the presented peptide.

According to the MS results, we used the method (56)constructed in our laboratory to score each amino acid in the peptide map and scanned the glycoproteins of 5 strains of MDV1 viruses: GA (GenBank accession no. AF147806), Md5 (GenBank accession no. NC_002229), RB1B (GenBank accession no. EF523390), 648A (GenBank accession no. JQ806361), and CVI988/Rispens (GenBank accession no. DQ530348). Finally, we screened 7 high-scoring nonapeptides that were the most conserved sequences in the 5 strains (Table 4), and three of the peptides are present in the eight originally predicted nonapeptides.

**Table 4.** Affinity prediction results obtained from the epitope scoring for virus-derived peptides.

| Derived from | Peptide | Potion | Score | CVI988 | GA | Md5 | RB1B | 648a |
| --- | --- | --- | --- | --- | --- | --- | --- | --- |
| Envelope glycoprotein E | IRYSRNGTR | 143-151 | 7.31 | + | + | + | + | + |
| Envelope glycoprotein I | RRIYIGEPR | 297-305 | 7.17 | + | + | + | + | + |
| Envelope glycoprotein E | RRRRRRRTR | 420-428 | 6.71 | + | + | + | + | + |
| MEQ<sup>*</sup> | RRRRKQTDY | 73-81 | 6.68 | - | + | + | + | + |
| MEQ<sup>*</sup> | RRRREQTDY | 73-81 | 6.68 | + | - | - | - | - |
| Envelope glycoprotein D<sup>*</sup> | RRRPSRRAY | 392-400 | 6.68 | + | + | + | + | + |
| Envelope glycoprotein H | IKFLNTGSY | 106-114 | 6.57 | + | + | + | + | + |

+ peptide exist in the strain
- peptide does not exist in the strain
<sup>\*</sup> represents the peptide present in the initial prediction Table S4

## Discussion

Based on the restriction between the TCR and MHC-I/II(57), SPF B15 chickens were selected to confirm the single epitope on tumorigenic MDV infection. First, the structural mechanism of the BF2*1501 molecule preferentially presenting MDV 9-mer peptide was clarified. Then, four CTL epitopes were identified by tetramer technique. In addition, each chicken was immunized with the dominant epitope, and the protective effect was objectively evaluated. The effect of single-epitope immunization, which can protect 33% of chickens from tumors, was determined for the first time, providing promising prospects for the continuous development of multiple epitope vaccines against MD and other oncogenic viruses.

To understand the functional mechanism of the BF2*1501 molecule presenting MDV epitopes, the crystal structures of BF2*1501 bound with the Meq-RY0801 and Meq-RY0901 epitopes were solved. The similarity of the two peptides is high, but the conformations of the Meq-RY0801 and Meq-RY0901 epitopes in the ABG are different: 8-mer-RY0801 is flatter, and 9-mer-RY0901 forms the “featured” M-type epitope morphology. Using rational analysis, these characteristics would make BF2*1501, which presents MDV 9-mer peptides, easier to be recognized by the TCR and may cause a more effective CTL immune response. In addition, pBF2*1501 has C, D and E pockets; these pockets have clear boundaries and relatively narrow volumes, which reduces the probability of presenting long peptides or long side chain amino acid peptides in the middle; the B and F pockets as anchor pockets are negatively charged; therefore, the P2 and P8/9 sites of the binding epitope peptide are non-negatively charged amino acid residues, which shows as a priority presentation feature of the BF2*1501 molecule, i.e., B15 chickens.

The candidate 9-mer epitopes from 5 MDV1 virulent strains were screened, and the results covered most of the functional proteins of MDV1. Finally, four candidate epitopes were selected from the gB and Meq proteins. Among them, 3 epitopes matched the gB protein of 5 strains, and 1 epitope matched only the vaccine strain and did not exist in other virulent strains. This result confirmed that the Meq mutation region may lead to immune escape. This result also limited the possibility of using gB plus Meq double epitopes in this experiment. Finally, three conserved epitopes were identified from the MDV gB protein. The GB0902 and GB0903 epitopes stimulated a stronger CTL immune response and are the dominant CTL epitopes. Therefore, we further clarified the immunoprotective effect of each epitope.

Human and mammalian gp96 belongs to the heat shock protein (HSP) 90 family, and the HSP90 family has a series of immunological functions(58). In particular, after binding peptides in vitro, the gp96-peptide complex is formed, which is delivered to the body and then released to MHC-I through endocytosis mediated by the receptor on the cell surface of APCs(59). The phagocytosis mediated by the gp96-peptide complex can induce a CTL immune response that is 10000 times higher than phagocytosis mediated by natural peptide presentation(60, 61). The peptide-binding region of gp96 is located in the N-terminus, and approximately 200 amino acids form an α-helix and several β-sheets(62). The peptide-binding groove of gp96 formed by a dimer is similar to the ABG of the pMHC-I complex (62). A large number of experiments have proven that the 333-aa N-terminal fragment of gp96 (excluding 22 signal peptides) is an excellent protein adjuvant for viral and tumor peptide vaccines(58, 63–68). Chicken HSP108 is a homolog of human and mammalian gp96(69). Its 333-aa N-terminal fragment is more than 90% homologous to human gp96, and its ability to be a protein adjuvant has been basically confirmed. In this research, chicken HSP-108-N333 was expressed and purified, combined with a single epitope as a protein adjuvant, mixed with IFA and injected into chickens. The preliminary and formal experimental results fully demonstrated its safety and efficiency. The number of CTLs increased by 9.3 times on average. We can be reasonably sure that without the HSP-108-N333 protein adjuvant, the challenge effect of single-epitope immunization is difficult to achieve 33%. Therefore, these results also showed the immunological potential of the HSP-108-N333 protein adjuvant for small-molecule epitope vaccines in chickens.

Compared with the MDV J-1 challenge group, the GB0902 and GB0903 epitopes provided 37.5% and 12.5% immune protection rates, respectively (Table 3). During the whole experiment, the mortality rate and tumorigenic rate of the J-1-challenge group were 40% and 80%, respectively. The mortality rate of the GB0902 epitope group decreased by 20%, and the tumorigenic rate was 50% (Table 3). The results confirmed that the GB0902 epitope could significantly reduce the mortality. It must be noted that the gold standard for the use of the MDV vaccine is that the immune protection rate should be above 80%. The gold standard evaluation of MD vaccines is to observe only whether there are tumors in viscera by the naked eye, but this paper used the results of tissue section staining and microscopic observation at the cytological level.

Safety and efficacy are the most important criteria of various vaccines. The epitope vaccine has no viral nucleic acid, so there is no viral replication or expelled virus. This is not only safe but can also alleviate the number of mutations of MDV strains. In this paper, the best single epitope, GB0902, with the protein adjuvant had an immune protection rate that reached 33%, which did not meet the requirements of the vaccine. For this reason, we further determined the antigen presentation profile of the B15 haplotype and searched for candidate epitopes for MDV strains. We further selected 7 candidate epitopes according to the antigen presentation profile and carried out a binding test in vitro. However, if an MDV epitope vaccine is used, it is necessary to identify MHC-I loci and MHC regions in individuals and populations of chickens. This is not only an advantage but also a disadvantage, especially when it is necessary to determine the MHC-I molecule before inoculating animal populations. The same is true for humans, and it is necessary to identify individual and family MHC-I loci to design CTL epitope vaccines accurately. Therefore, we speculate that the immune effect of multiepitope vaccines will reach the gold standard, and the addition of B- and Th-cell epitopes may become a “super epitope vaccine”.

In conclusion, this study confirmed that the GB0902 epitope is the dominant CTL epitope, and the protection rate against the vvMDV J-1 strain is 33%, which clearly shows the safety and effectiveness of a single epitope. We believe that with the further development of T cell and B cell epitopes, the “super epitope vaccine” is able to have a newer and more ideal effect in the field of oncogenic virus control in jawed vertebrates.

## Materials and Methods

### Animals, virus, and peptides

SPF White Leghorn chickens were provided by Merial (now Boehringer-Ingelheim). This breed is a B15 haplotype; that is, it is a purebred lineage with the BF2*1501 allele (expressing BF2*1501). After the chickens hatched in the SPF chicken house, they were kept in a positive- and negative-pressure poultry isolator for experiments. The vvMDV J-1 strain was provided by the China Institute of Veterinary Drugs Control. The peptides used in this experiment are listed in S3 Table. These peptides and the synthetic random peptide library Ran_9Xsplitted (9X, XXXXXXXXX, where X is a random amino acid other than cysteine) used in this study were synthesized and purified by reverse-phase high-performance liquid chromatography (HPLC, SciLight Biotechnology, Beijing) with >90% purity. The lyophilized peptides were stored at −80°C and dissolved in dimethyl sulfoxide at a suitable concentration before use.

### Protein preparation

Recombinant expression plasmids containing the BF2*1501 heavy chain (HC) gene (GenBank accession no. L28958) and chicken β2m gene (GenBank accession no. BAD22696) were constructed previously(70). The constructed plasmids were transformed into the *E. coli* BL21(DE3) strain for prokaryotic expression to obtain inclusion bodies, and then the proteins were purified as previously described(71). These purified proteins were dissolved in 6 M guanidine hydrochloride at a final concentration of 30 mg/ml and stored at −20°C.

### The assembly and purification of the pBF2*1501 complexes

The protocol was essentially the same as that described previously(71). The BF2*1501 HC and chicken β2m inclusion bodies and a peptide were refolded at a 1:1:1 molar ratio. After incubation for 24 hours at 4°C, the soluble portion was concentrated and purified by chromatography on a Superdex 200 16/60 HiLoad size-exclusion column (GE Healthcare) followed by a Resource Q anion-exchange chromatography column (GE Healthcare). The purified protein was finally dialysed against buffer (20 mM Tris-HCl (pH 8.0) and 50 mM NaCl).

### Crystallization of pBF2*1501 and data collection

The purified complex (45 kDa) of pBF2*1501 with an MDV-derived peptide was dialyzed against crystallization buffer (20 mM Tris–HCl (pH 8.0) and 50 mM NaCl) and concentrated to 12 mg/ml. The commercially available Index kit (Hampton Research, California, USA) consisting of preformulated solutions was used for crystallization. In total, 1 μl of protein solution and 1 μl of reservoir crystallization buffer were placed over a well containing 160 ml of reservoir solution using a VDX plate (Hampton Research) at 291 K. Crystals were obtained under optimized conditions (0.1 M Tris–HCl (pH 8.5) and 1.9 M ammonium sulfate at 291 K). The crystals were soaked for 20-30 s in reservoir solution supplemented with 20% glycerol as a cryoprotectant and then directly flash-cooled in liquid nitrogen. Diffraction data sets were collected at the Shanghai Synchrotron Radiation Facility (SSRF) using beamline BL17U at a wavelength of 0.97916 Å (Shanghai, China). The collected intensities were indexed, integrated, corrected for absorption, scaled and merged using HKL2000(72).

### Structure determination, refinement, and data analysis

The structure of BF2*1501 with the Meq-RY0801 or Meq-RY0901 peptide was determined through the molecular replacement method with the PHASER program in CCP4i with BF2*2101 (PDB ID: 3BEV) as the search model. After correction of the maps in Coot and several rounds of refinement in REFMAC5, phenix.refine was used for further refinement. Finally, MolProbity tools were used in Phenix for model quality assessment. The structural illustrations and the related figures were generated by PyMOL v0.99rc2 (Schrödinger, LLC). Sequence alignment was performed by Clustal Omega (https://www.ebi.ac.uk/Tools/msa/clustalo/) and ESPript 3.0 (http://espript.ibcp.fr/ESPript/ESPript/). Accessible surface area (ASA) and buried surface area (BSA) values were calculated by the online PDBePISA website (http://www.ebi.ac.uk/msd-srv/prot_int/pistart.html).

### Preparation and identification of epitope-specific tetramers

The recombinant expression plasmid pET-21a(+)/BF2*1501-BSP incorporated BF2*1501 HC with the BirA substrate peptide (BSP), which is necessary for tetramerization and was constructed previously(70). The expression, assembly with the GB0902, GB0903, GB0905 and RY0903 peptides, and purification of pBF2*1501-BSP were the same as described above. After purification, the complex was further biotinylated using the BirA enzyme (Epigen Biosciences, Ltd., Beijing, China) and then purified by chromatography on a Superdex 200 16/60 HiLoad size-exclusion column (GE Healthcare). Finally, pBF2*1501 epitope-specific tetramers were prepared by mixing the biotinylated pBF2*1501 complexes and phycoerythrin (PE)-labeled streptavidin (Sigma Aldrich) at a molar ratio of 4:1, after which the complexes were separated using a 100-kDa Millipore filter. SDS-PAGE was used to evaluate the efficiency of tetramerization.

### Identification of dominant CTL epitopes of MDV

A total of six one-day-old SPF chickens expressing BF2*1501 molecules were provided by Merial (now Boehringer-Ingelheim) in Beijing and divided randomly into two groups: the vaccine-immunized group and the control group (3 chickens in each group). Chickens in the vaccine-immunized group were initially immunized with the commercially available MDV live-attenuated CVI988/Rispens vaccine (Merial, now Boehringer-Ingelheim) at 1 day of age, according to the manufacturer’s instructions. After seven days, the immunized chickens were subcutaneously injected with the incomplete Freund’s adjuvant (IFA)-emulsified (1:3 emulsification) GB0902, GB0903, GB0905 and RY0903 peptides (0.5 mg/kg bodyweight). Chickens in the control group were treated with PBS. After 14 days, peripheral blood mononuclear cells (PBMCs) were isolated using Ficoll-Paque according to the manufacturer’s instructions (Haoyang Biological Manufacture Company, Tianjin, China). PBMCs were incubated with fluorescence-activated cell sorting (FACS) buffer (PBS with 0.1% bovine serum albumin and 0.1% sodium azide) containing the PE-labeled tetrameric pBF2*1501 complex and fluorescein isothiocyanate (FITC)-labeled anti-CD8 monoclonal antibody (SouthernBiotech, USA) for 30 min at 4°C. Subsequently, the PBMCs were washed twice with FACS buffer and immobilized with 1% paraformaldehyde. Flow cytometry was used to determine the MDV CTL epitope. More than 10^6^ cell events were acquired for each sample. Cells that were positive for both PE-labeled pBF2*1501 tetramers and the FITC-labeled anti-CD8 monoclonal antibody were counted as epitope-responsive CD8^+^ T cells. GraphPad Prism was used to analyze the data. Significant differences (P<0.05) between means were assessed by a two-tailed Student’s t-test(70).

### Preparation and identification of the protein adjuvant HSP108-N333

Total RNA was extracted from chicken liver cells for reverse transcription, and the HSP108-N333 gene (encoding the N-terminal fragment (23-355 amino acid (aa)), GenBank accession no. NM_204289) was amplified using cDNA as a template (forward: 5’-CGGGATCCGATGATGAAGTTGTTCAGCGTGAGG-3’; reverse: 5’-CCGCTCGAGTTAAGCAGTAAAATGGATGTAAGCCAT-3’). PCR conditions were: 98°C, 5 min; 98°C, 30 s; 55°C, 30 s; 72°C, 1 min; 30 cycle; and 72°C, 10 min. The gene was inserted into the pGEX-6P-1 vector and named pGEX-6P-1/HSP108-N333(68). The fusion protein HSP108-N333-Glutathione S-transferase (GST) was obtained by IPTG-induced protein expression, and the GST tag was removed by PreScission Protease (PSP) digestion. The HSP108-N333 protein monomer was purified by a HiLoad 16/60 Superdex 200 gel filtration column and identified by SDS-PAGE. Subsequently, 100 µg of HSP108-N333 protein and 100 µg of CTL epitope peptide GB0903 were combined in binding buffer (20 mM Tris-Cl (pH 8.0), 20 mM NaCl, 2 mM MgCl_2_, and 120 mM KCl) at 37°C for 20 min, cooled to room temperature, and then mixed with Freund’s adjuvant at a 1:1 volume ratio to immunize SPF chickens. A total of 3 immunizations were performed one week apart each time. One week after the third immunization, PBMCs were separated, stained with the FITC-labeled anti-chicken CD8 monoclonal antibody and PE-labeled BF2*1501-BSP/GB0903 tetramer, and analyzed by flow cytometry.

### The immunoprotective effect of epitopes

A total of forty 1-day-old SPF chickens were divided randomly into four groups: the control group, the challenge group, the GB0902 epitope-immunized (GB0902 epitope) group and the GB0903 epitope-immunized (GB0902 epitope) group (10 chickens in each group). The GB0902 and GB0903 epitope groups were subcutaneously injected with the peptide vaccine, which included the protein adjuvant HSP108-N333 (0.5 mg/kg) at 1 and 7 days of age, and the vaccine preparation method was the same as above. Each chicken in the control and challenge groups was subcutaneously injected with PBS at 1 and 7 days of age. On the fifth day after the second immunization, the vvMDV J-1 strain (10^2^ LD_50_) was diluted 10-fold, and 0.2 ml was injected into each chicken of the GB0902 epitope group, the GB0903 epitope group, and the challenge group. The observation period lasted for 42 days. The morbidity and mortality of each group were observed and recorded every day, and the dead chickens were dissected to observe the pathological changes. The criteria for MD-positive chickens were as follows: the chickens that died during the experiment were determined to have MD based on the clinical symptoms and visual lesions. On the last day of the experiment, all surviving chickens were sacrificed and dissected, and those with visual tumors were determined to be MD-positive. At the end of the experiment, the liver, kidney and spleen of all chickens were collected for paraffin sectioning, and hematoxylin-eosin staining was used for pathological observation. The PI and disease index (DI) were calculated according to the results.

The immune efficacy of the experimental vaccine was expressed by the PI:

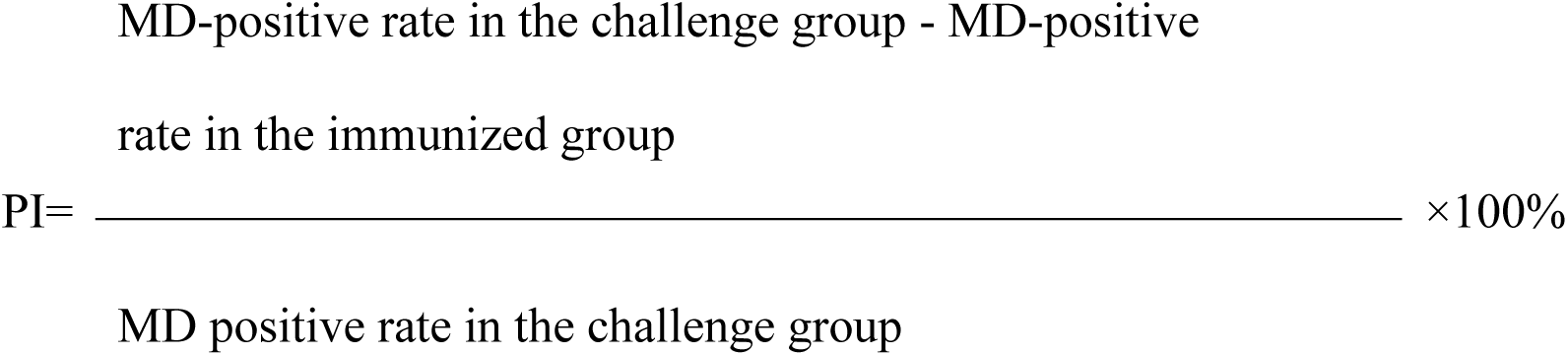

The pathological changes in the immunized chickens were expressed by the DI:

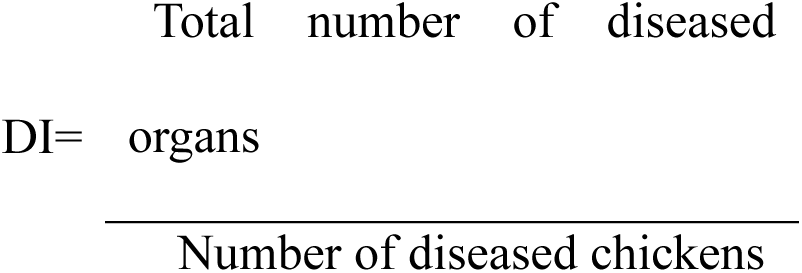

### Liquid chromatography–tandem mass spectrometry

pBF2*1501 refolded with the Ran_9Xsplitted library was treated with 0.2 N acetic acid, incubated at 65°C for 30 min, and then concentrated with a 3 kDa filter to collect the peptides. The samples were desalinated by a method previously described(73). Peptide motifs of the single dominantly expressed class I molecule explain the striking MHC-determined response to Rous sarcoma virus in chickens. First, 200 μl of methanol was used to activate the desalting tips, and 200 μl of 0.1% (v/v) trifluoroacetic acid (TFA) was used to equilibrate the desalting tips. Then, the peptides were washed twice with 200 μl of washing buffer (0.1% [v/v] TFA) and eluted with 200 μl of eluting solution (0.1% [v/v] TFA and 75% [v/v] acetonitrile). The eluted peptides were lyophilized and stored at −80°C. The peptides were separated with the Easy nano-LC1000 system (Thermo Fisher Scientific, San Jose, CA) as previously described. The acquisition of mass spectrometry (MS) data was carried out by a Q Exactive HF system (Thermo Fisher Scientific, Bremen, Germany) in the data-dependent acquisition mode, and the top 20 precursors by intensity with a mass range of m/z 300 to 1800 were sequentially fragmented with higher-energy collisional dissociation and normalized collision energy of 27. The dynamic exclusion time was 20 s. Automatic gain control for MS1 and MS2 was set to 3e^6^ and 1e, and the resolution for MS1 and MS2 was set to 120 and 30K, respectively.

### Analysis of de novo and candidate epitope scoring

Based on the peptide-binding profile information, the detection threshold was adjusted (sequence length = 9 and score ≥ 90). In addition, to determine the restricted motif of pBF2*1501 presentation, the standard deviation (σ), average value (^x̅^) and coefficient of variation (Vs) were calculated as previously described(56). A peptide position with a high Vs value was predicted to be a restricted position in the presentation. Peptides with a high affinity to BF2*1501 were predicted according to a previous method(56).

### Protein structure accession number

The structures of pBF2*1501 with the MDV-RY0801 peptide and pBF2*1501 with the MDV-RY0901 peptide were deposited in the PDB (http://www.rcsb.org/pdb/home/home.do) with the PDB ID of 7DBY, 7DBZ, respectively.

### Ethics statement

The animal trials in this study were performed according to the Chinese Regulations for Laboratory Animals-The Guidelines for the Care of Laboratory Animals (Ministry of Science and Technology of People’s Republic of China) and Laboratory Animal Requirements for Environment and Housing Facilities (GB14925-2010, National Laboratory Animal Standardization Technical Committee). The license number associated with this research protocol is CAU20140305-2, which was approved by The Laboratory Animal Ethics Committee of China Agricultural University. The protocol adhered to the recommendations in the Institute for Laboratory Animals Research’s “Guide for the Care and Use of Laboratory Animals”.

## Acknowledgments

This work was supported financially by grants from the National Natural Science Foundation of China (NSFC) (grant no.31972683 and 31572493, http://www.nsfc.gov.cn). We thank the Shanghai Synchrotron Radiation Facility (SSRF) for the crystal diffraction data and everyone who has assisted in this work.

## Supporting Information

**S1 Table. The interactions between the peptide and the ABG of pBF2*1501.**

**S2 Table. The ASA and BSA calculations of the bound peptide.**

**S3 Table. Refolding of BF2*1501 and chicken β2m with MDV-derived peptides.**

**S4 Table. Results of LC-MS/MS de novo analysis.**

## Reference

1. Haq K, Schat KA, Sharif S. 2013. Immunity to Marek’s disease: where are we now? Dev Comp Immunol 41:439–46.

2. Bailey RI, Cheng HH, Chase-Topping M, Mays JK, Anacleto O, Dunn JR, Doeschl-Wilson A. 2020. Pathogen transmission from vaccinated hosts can cause dose-dependent reduction in virulence. PLoS Biol 18:e3000619.

3. Gimeno IM, Schat KA. 2018. Virus-Induced Immunosuppression in Chickens. Avian Dis 62:272–285.

4. Rozins C, Day T, Greenhalgh S. 2019. Managing Marek’s disease in the egg industry. Epidemics 27:52–58.

5. Fukuchi K, Sudo M, Lee YS, Tanaka A, Nonoyama M. 1984. Structure of Marek’s disease virus DNA: detailed restriction enzyme map. J Virol 51:102–9.

6. Tulman ER, Afonso CL, Lu Z, Zsak L, Rock DL, Kutish GF. 2000. The genome of a very virulent Marek’s disease virus. J Virol 74:7980–8.

7. Lee L, Wu P, Sui D, Ren D, Kamil J, Kung H, Witter R. 2000. The complete unique long sequence and the overall genomic organization of the GA strain of Marek’s disease virus. Proceedings of the National Academy of Sciences 97.

8. Calnek BW. 2001. Pathogenesis of Marek’s disease virus infection. Curr Top Microbiol Immunol 255:25–55.

9. Baaten BJ, Staines KA, Smith LP, Skinner H, Davison TF, Butter C. 2009. Early replication in pulmonary B cells after infection with Marek’s disease herpesvirus by the respiratory route. Viral Immunol 22:431–44.

10. Chakraborty P, Vervelde L, Dalziel RG, Wasson PS, Nair V, Dutia BM, Kaiser P. 2017. Marek’s disease virus infection of phagocytes: a de novo in vitro infection model. J Gen Virol 98:1080–1088.

11. Levy AM, Gilad O, Xia L, Izumiya Y, Choi J, Tsalenko A, Yakhini Z, Witter R, Lee L, Cardona CJ, Kung HJ. 2005. Marek’s disease virus Meq transforms chicken cells via the v-Jun transcriptional cascade: a converging transforming pathway for avian oncoviruses. Proc Natl Acad Sci U S A 102:14831–6.

12. Brown AC, Baigent SJ, Smith LP, Chattoo JP, Petherbridge LJ, Hawes P, Allday MJ, Nair V. 2006. Interaction of MEQ protein and C-terminal-bindingprotein is critical for induction of lymphomas by Marek’s disease virus. Proc Natl Acad Sci U S A 103:1687–92.

13. Rozins C, Day T. 2017. The industrialization of farming may be driving virulence evolution. Evol Appl 10:189–198.

14. Read AF, Baigent SJ, Powers C, Kgosana LB, Blackwell L, Smith LP, Kennedy DA, Walkden-Brown SW, Nair VK. 2015. Imperfect Vaccination Can Enhance the Transmission of Highly Virulent Pathogens. PLoS Biol 13:e1002198.

15. Osterrieder NV, J.F. 2004. The genome content of Marek’s disease-like viruses. Curr Top Microbiol.

16. Bertzbach LD, Pfaff F, Pauker VI, Kheimar AM, Höper D, Härtle S, Karger A, Kaufer BB. 2019. The Transcriptional Landscape of Marek’s Disease Virus in Primary Chicken B Cells Reveals Novel Splice Variants and Genes. Viruses 11.

17. Tischer BK, Schumacher D, Beer M, Beyer J, Teifke JP, Osterrieder K, Wink K, Zelnik V, Fehler F, Osterrieder N. 2002. A DNA vaccine containing an infectious Marek’s disease virus genome can confer protection against tumorigenic Marek’s disease in chickens. J Gen Virol 83:2367–2376.

18. Yang Y, Dong M, Hao X, Qin A, Shang S. 2020. Revisiting cellular immune response to oncogenic Marek’s disease virus: the rising of avian T-cell immunity. Cell Mol Life Sci 77:3103–3116.

19. Osterrieder N, Kamil JP, Schumacher D, Tischer BK, Trapp S. 2006. Marek’s disease virus: from miasma to model. Nat Rev Microbiol 4:283–94.

20. Paulette F. Suchodolski YI, Blanca Lupiani, Dharani K. Ajithdoss, Lucy F. Lee, Hsing-Jien Kung, Sanjay M. Reddy. 2010. Both homo and heterodimers of Marek’s disease virus encoded Meq protein contribute to transformation of lymphocytes in chickens. Virology 399:312–21.

21. Lee LF, Lupiani B, Silva RF, Kung HJ, Reddy SM. 2008. Recombinant Marek’s disease virus (MDV) lacking the Meq oncogene confers protection against challenge with a very virulent plus strain of MDV. Vaccine 26:1887–92.

22. Schat KA, Xing Z. 2000. Specific and nonspecific immune responses to Marek’s disease virus. Dev Comp Immunol 24:201–21.

23. Nair V. 2013. Latency and tumorigenesis in Marek’s disease. Avian Dis 57:360–5.

24. Witter RL. 1997. Increased virulence of Marek’s disease virus field isolates. Avian Dis 41:149–63.

25. Faiz NM, Cortes AL, Guy JS, Fletcher OJ, Cimino T, Gimeno IM. 2017. Evaluation of factors influencing the development of late Marek’s disease virus-induced immunosuppression: virus pathotype and host sex. Avian Pathol 46:376–385.

26. Witter RL, Gimeno IM, Reed WM, Bacon LD. 1999. An acute form of transient paralysis induced by highly virulent strains of Marek’s disease virus. Avian Dis 43:704–20.

27. Gimeno IM, Witter RL, Reed WM. 1999. Four distinct neurologic syndromes in Marek’s disease: effect of viral strain and pathotype. Avian Dis 43:721–37.

28. Gimeno IM, Witter RL, Hunt HD, Lee LF, Reddy SM, Neumann U. 2001. Marek’s disease virus infection in the brain: virus replication, cellular infiltration, and major histocompatibility complex antigen expression. Vet Pathol 38:491–503.

29. Witter RL, Gimeno IM. 2006. Susceptibility of adult chickens, with and without prior vaccination, to challenge with Marek’s disease virus. Avian Dis 50:354–65.

30. Dunn JR, Black Pyrkosz A, Steep A, Cheng HH. 2019. Identification of Marek’s disease virus genes associated with virulence of US strains. J Gen Virol 100:1132–1139.

31. Schat KA. 2016. History of the First-Generation Marek’s Disease Vaccines: The Science and Little-Known Facts. Avian Dis 60:715–724.

32. Reddy SM, Izumiya Y, Lupiani B. 2017. Marek’s disease vaccines: Current status, and strategies for improvement and development of vector vaccines. Vet Microbiol 206:113–120.

33. Nair V. 2018. Hot Topics in Avian Pathology: Marek’s disease. Avian Pathology.

34. Singh SM, Baigent SJ, Petherbridge LJ, Smith LP, Nair VK. 2010. Comparative efficacy of BAC-derived recombinant SB-1 vaccine and the parent wild type strain in preventing replication, shedding and disease induced by virulent Marek’s disease virus. Res Vet Sci 89:140–5.

35. Atkins KE, Read AF, Savill NJ, Renz KG, Walkden-Brown SW, Woolhouse ME. 2011. Modelling Marek’s disease virus (MDV) infection: parameter estimates for mortality rate and infectiousness. BMC Vet Res 7:70.

36. Davison F, Nair V. 2005. Use of Marek’s disease vaccines: could they be driving the virus to increasing virulence? Expert Rev Vaccines 4:77–88.

37. Gimeno IM. 2008. Marek’s disease vaccines: a solution for today but a worry for tomorrow? Vaccine 26 Suppl 3:C31–41.

38. Xinyu Zhuang HZ, Huoying Shi, Hongxia Shao, Jianqiang Ye, Ji Miao, Genghua Wu and Aijian Qin*. 2015. Outbreak of Marek’s disease in a vaccinated broiler breeding flock during its peak egg-laying period in China. BMC Vet Res 11:157.

39. Muniyellappa HK, Satyanarayana ML, Isloor S, Shivakumar Gowda NK. 2013. Marek’s disease outbreak among vaccinated commercial layer flocks in the mining area of Karnataka, India. Vet Rec 172:452.

40. Lee LE, Witter RL, Reddy SM, Wu P, Yanagida N, Yoshida S. 2003. Protection and synergism by recombinant fowl pox vaccines expressing multiple genes from Marek’s disease virus. Avian Dis 47:549–58.

41. Lee LF, Kreager KS, Arango J, Paraguassu A, Beckman B, Zhang H, Fadly A, Lupiani B, Reddy SM. 2010. Comparative evaluation of vaccine efficacy of recombinant Marek’s disease virus vaccine lacking Meq oncogene in commercial chickens. Vaccine 28:1294–9.

42. Su S, Cui N, Li J, Sun P, Li H, Li Y, Cui Z. 2016. Deletion of the BAC sequences from recombinant meq-null Marek’s disease (MD) virus increases immunosuppression while maintaining protective efficacy against MD. Poult Sci 95:1504–1512.

43. Zhang Y, Tang N, Sadigh Y, Baigent S, Shen Z, Nair V, Yao Y. 2018. Application of CRISPR/Cas9 Gene Editing System on MDV-1 Genome for the Study of Gene Function. Viruses 10.

44. Weidang Li MDJ, Smita Singhania, Kyle H. Ramsey and Ashlesh K. Murthy. 2014. Peptide Vaccine: Progress and Challenges. Vaccines (Basel) 2:515–36.

45. Bartnik A, Nirmal AJ, Yang SY. 2012. Peptide Vaccine Therapy in Colorectal Cancer. Vaccines (Basel) 1:1–16.

46. Yoshihiro Oka AT, Jun Nakata, Sumiyuki Nishida, Naoki Hosen, Atsushi Kumanogoh, Yusuke Oji, Haruo Sugiyama. 2017. Wilms’ Tumor Gene 1 (WT1) Peptide Vaccine Therapy for Hematological Malignancies: From CTL Epitope Identification to Recent Progress in Clinical Studies Including a Cure-Oriented Strategy. Oncol Res Treat 40:682–690.

47. Baba T, Sato-Matsushita M, Kanamoto A, Itoh A, Oyaizu N, Inoue Y, Kawakami Y, Tahara H. 2010. Phase I clinical trial of the vaccination for the patients with metastatic melanoma using gp100-derived epitope peptide restricted to HLA-A*2402. J Transl Med 8:84.

48. Mittendorf EA, Holmes JP, Ponniah S, Peoples GE. 2008. The E75 HER2/neu peptide vaccine. Cancer Immunol Immunother 57:1511–21.

49. Abdulla F, Adhikari UK, Uddin MK. 2019. Exploring T & B-cell epitopes and designing multi-epitope subunit vaccine targeting integration step of HIV-1 lifecycle using immunoinformatics approach. Microb Pathog 137:103791.

50. Leung HC, Chan CC, Poon VK, Zhao HJ, Cheung CY, Ng F, Huang JD, Zheng BJ. 2015. An H5N1-based matrix protein 2 ectodomain tetrameric peptide vaccine provides cross-protection against lethal infection with H7N9 influenza virus. Emerg Microbes Infect 4:e22.

51. & ZZLPYDPZJLHC, & YFXLHCJZJSTL, Wang FZYZY. 2015. Efficacy of synthetic peptide candidate vaccines against serotype-A foot-and-mouth disease virus in cattle. Appl Microbiol Biotechnol 99:1389–98.

52. Marrack P, Scott-Browne JP, Dai S, Gapin L, Kappler JW. 2008. Evolutionarily conserved amino acids that control TCR-MHC interaction. Annu Rev Immunol 26:171–203.

53. Schat KA, Taylor RL, Jr., Briles WE. 1994. Resistance to Marek’s disease in chickens with recombinant haplotypes to the major histocompatibility (B) complex. Poult Sci 73:502–8.

54. Koch M, Camp S, Collen T, Avila D, Salomonsen J, Wallny HJ, van Hateren A, Hunt L, Jacob JP, Johnston F, Marston DA, Shaw I, Dunbar PR, Cerundolo V, Jones EY, Kaufman J. 2007. Structures of an MHC class I molecule from B21 chickens illustrate promiscuous peptide binding. Immunity 27:885–99.

55. Wallny HJ, Avila D, Hunt LG, Powell TJ, Riegert P, Salomonsen J, Skjødt K, Vainio O, Vilbois F, Wiles MV, Kaufman J. 2006. Peptide motifs of the single dominantly expressed class I molecule explain the striking MHC-determined response to Rous sarcoma virus in chickens. Proc Natl Acad Sci U S A 103:1434–9.

56. Qu Z, Li Z, Ma L, Wei X, Zhang L, Liang R, Meng G, Zhang N, Xia C. 2019. Structure and Peptidome of the Bat MHC Class I Molecule Reveal a Novel Mechanism Leading to High-Affinity Peptide Binding. The Journal of Immunology 202:3493–3506.

57. La Gruta NL, Gras S, Daley SR, Thomas PG, Rossjohn J. 2018. Understanding the drivers of MHC restriction of T cell receptors. Nat Rev Immunol 18:467–478.

58. Suto R, Srivastava PK. 1995. A mechanism for the specific immunogenicity of heat shock protein-chaperoned peptides. Science 269:1585–8.

59. Berwin B, Rosser MF, Brinker KG, Nicchitta CV. 2002. Transfer of GRP94(Gp96)-associated peptides onto endosomal MHC class I molecules. Traffic 3:358–66.

60. Bolhassani A, Rafati S. 2008. Heat-shock proteins as powerful weapons in vaccine development. Expert Rev Vaccines 7:1185–99.

61. Arnold-Schild D, Hanau D, Spehner D, Schmid C, Rammensee HG, de la Salle H, Schild H. 1999. Cutting edge: receptor-mediated endocytosis of heat shock proteins by professional antigen-presenting cells. J Immunol 162:3757–60.

62. Linderoth NA, Popowicz A, Sastry S. 2000. Identification of the peptide-binding site in the heat shock chaperone/tumor rejection antigen gp96 (Grp94). J Biol Chem 275:5472–7.

63. Liu B, Dai J, Zheng H, Stoilova D, Sun S, Li Z. 2003. Cell surface expression of an endoplasmic reticulum resident heat shock protein gp96 triggers MyD88-dependent systemic autoimmune diseases. Proc Natl Acad Sci U S A 100:15824–9.

64. Zugel U, Sponaas AM, Neckermann J, Schoel B, Kaufmann SH. 2001. gp96-peptide vaccination of mice against intracellular bacteria. Infect Immun 69:4164–7.

65. Li HT, Yan JB, Li J, Zhou MH, Zhu XD, Zhang YX, Tien P. 2005. Enhancement of humoral immune responses to HBsAg by heat shock protein gp96 and its N-terminal fragment in mice. World J Gastroenterol 11:2858–63.

66. Li Z, Dai J, Zheng H, Liu B, Caudill M. 2002. An integrated view of the roles and mechanisms of heat shock protein gp96-peptide complex in eliciting immune response. Front Biosci 7:d731–51.

67. Liu Z, Li X, Qiu L, Zhang X, Chen L, Cao S, Wang F, Meng S. 2009. Treg suppress CTL responses upon immunization with HSP gp96. Eur J Immunol 39:3110–20.

68. Li H, Zhou M, Han J, Zhu X, Dong T, Gao GF, Tien P. 2005. Generation of murine CTL by a hepatitis B virus-specific peptide and evaluation of the adjuvant effect of heat shock protein glycoprotein 96 and its terminal fragments. J Immunol 174:195–204.

69. Poola I, Kiang JG. 1994. The estrogen-inducible transferrin receptor-like membrane glycoprotein is related to stress-regulated proteins. J Biol Chem 269:21762–9.

70. Li X, Zhang L, Liu Y, Ma L, Zhang N, Xia C. 2020. Structures of the MHC-I molecule BF2*1501 disclose the preferred presentation of an H5N1 virus-derived epitope. The Journal of biological chemistry doi:10.1074/jbc.RA120.012713.

71. Sun B, Li X, Wang Z, Xia C. 2013. Complex assembly, crystallization and preliminary X-ray crystallographic analysis of the chicken MHC class I molecule BF2*1501. Acta Crystallogr Sect F Struct Biol Cryst Commun 69:122–5.

72. Jensen LH. 1997. Refinement and reliability of macromolecular models based on X-ray diffraction data. Methods Enzymol 277:353–66.

73. Ma L, Zhang N, Qu Z, Liang R, Zhang L, Zhang B, Meng G, Dijkstra JM, Li S, Xia MC. 2020. A Glimpse of the Peptide Profile Presentation by Xenopus laevis MHC Class I: Crystal Structure of pXela-UAA Reveals a Distinct Peptide-Binding Groove. The Journal of Immunology 204:147–158.

